# Coral species-specific loss and physiological legacy effects are elicited by extended marine heatwave

**DOI:** 10.1101/2023.09.18.558296

**Authors:** E.L. Strand, K.H. Wong, A. Farraj, S. Gray, A. McMenamin, H.M. Putnam

## Abstract

Marine heatwaves are increasing in frequency and intensity, with potentially catastrophic consequences for marine ecosystems such as coral reefs. An extended heatwave and recovery time-series that incorporates multiple stressors and is environmentally realistic can provide enhanced predictive capacity for performance under climate change conditions. We exposed common reef-building corals in Hawai‘i, *Montipora capitata* and *Pocillopora acuta*, to a 2-month period of high temperature and high pCO_2_ conditions or ambient conditions in a factorial design, followed by 2 months of ambient conditions. High temperature, rather than high pCO_2_, drove multivariate physiology shifts through time in both species, including decreases in respiration rates and endosymbiont densities. *Pocillopora acuta* exhibited more significantly negatively altered physiology, and substantially higher bleaching and mortality than *M. capitata*. The sensitivity of *P. acuta* appears to be driven by higher baseline rates of photosynthesis paired with lower host antioxidant capacity, creating an increased sensitivity to oxidative stress. Thermal tolerance of *M. capitata* may be partly due to harboring a mixture of *Cladocopium* and *Durusdinium* spp., whereas *P. acuta* was dominated by other distinct *Cladocopium* spp. Only *M. capitata* survived the experiment, but physiological state in heatwave exposed *M. capitata* remained significantly diverged at the end of recovery relative to individuals that experienced ambient conditions. In future climate scenarios, particularly marine heatwaves, our results indicate a species-specific loss of corals that is driven by baseline host and symbiont physiological differences as well as Symbiodiniaceae community compositions, with the surviving species experiencing physiological legacies that are likely to influence future stress responses.

## Introduction

Coral holobionts, a meta-organism composed of an animal coral host, endosymbiont dinoflagellates and associated microbial community (Rohwer et al., 2002), make up the structure of one of the most ecologically diverse and economically valuable ecosystems in the world: coral reefs. The function and health of the coral holobiont, and therefore coral reefs, is dependent on this multipartner symbiotic relationship that cycles nutrients and provides each partner with otherwise limited or unavailable nutritional resources (Bourne et al., 2016). However, climate change is impacting the marine environment through increasing ocean temperatures (IPCC, 2018), specifically marine heatwaves (Oliver et al., 2021) and ocean acidification (Gattuso et al., 2015), among other local stressors, challenging the symbiotic function, physiological processes and, ultimately, the survival of reef-building corals. Although ocean acidification will place a chronic energetic strain on coral calcification (Mollica et al., 2018) and cellular acid–base balance (Tresguerres et al., 2017; Gibbin et al., 2014), the dominant physiological stress is currently in the form of increasingly frequent, intense marine heatwaves (Oliver et al., 2018; Sweet and Brown, 2016). This thermal stress triggers dysbiosis known as ‘coral bleaching’, or the disruption of the symbiotic relationship between a coral host and their photosynthetic endosymbionts, Symbodinicaeae. This results in the coral host expelling the endosymbionts, leaving a stark white (‘bleached’) appearance and the coral host depleted of their primary carbon source (Gates et al., 1992; Boilard et al., 2020). Mass coral bleaching events resulting from marine heatwaves have increased in frequency and intensity in recent years (Oliver et al., 2021), and this trend is on track to continue, with catastrophic consequences projected for coral reef ecosystems (Heron et al., 2016; Le Nohaïc et al., 2017; van Woesik and Kratochwill, 2022).

The current understanding of the mechanistic cascade of bleaching is linked to the response of the host and/or endosymbionts to increased reactive oxygen species (ROS) levels produced under higher temperature that consequently increase oxidative stress and cause a breakdown of photosynthetic processes within the symbiont cells (Weis, 2008; Gates et al., 1992). This breakdown causes the symbiont cells to be either digested or expelled via several mechanisms including exocytosis, apoptosis, necrosis and host cell detachment (Weis, 2008; Lesser, 2004; Gates et al., 1992). A review of recent studies has shown that oxidative stress can occur in the host first (Oakley and Davy, 2018), suggesting that host antioxidant capacity may be critical in tolerating higher levels of temperature stress. However, host response to changing environment conditions has been shown to be highly species-specific (Bahr et al., 2016). In Kāne‘ohe Bay, O’ahu, Hawai’i, two common reef-building coral species display differential thermal tolerance: *Montipora capitata* is more resistant and *Pocillopora acuta* is more sensitive to environmental stress (Bahr et al., 2015; Gibbin et al., 2015). The underlying mechanisms to differential tolerance are known to be complex, ranging from genetic influences, epigenetic patterns, and associated endosymbiont and bacterial communities (Putnam, 2021), to host morphological structure and tissue thickness differences (Loya et al., 2001), but are not fully understood. Thus, multivariate analyses of key processes and traits can provide the capacity to quantify temporal variation in host, symbiont and holobiont responses, and are necessary to more fully elucidate the bleaching cascade (Gardner et al., 2017; Wall et al., 2021; McLachlan et al., 2021).

Although extensive research has been conducted on the effects of increased temperature on coral (Mydlarz et al., 2010; Sweet and Brown, 2016; McLachlan et al., 2020; Cziesielski et al., 2019; Lesser, 2011; van Oppen and Lough, 2018), there are fewer experimental studies that simultaneously address critical environmentally relevant aspects that reflect a coral’s natural conditions. Specifically, there are few studies that concurrently: (1) mimic daily and seasonal environmental fluctuations (van Woesik et al., 2022; Ziegler et al., 2021; Putnam and Edmunds, 2011), while (2) accounting for simultaneous, multivariate stressors (Pendleton et al., 2016; van Woesik et al., 2022), (3) sampling with high frequency to capture short and long-term temporal stress and recovery dynamics (Claar et al., 2020; Gardner et al., 2017), (4) during environmentally realistic times of the year (vanWoesik et al., 2022; Ziegler et al., 2021) and (5) tracking the survivors following the stress exposure (Claar et al., 2020; Gardner et al., 2017). These factors are particularly important, as exposure to diurnal temperature and pCO_2_ fluctuations elicit different responses than stable, or less variable conditions (Putnam and Edmunds, 2011; Dufault et al., 2012; Schoepf et al., 2022; Barshis et al., 2013; Wall et al., 2021). Further, high-frequency sampling of a variety of variables was able to elucidate that symbiont expulsion and bleaching precedes severe holobiont physiological responses and health decline in later stress time points (Gardner et al., 2017). Additionally, seasonal studies of influences on coral physiology (Jurriaans and Hoogenboom, 2020; Scheufen et al., 2017; Thornhill et al., 2011; Edmunds and Putnam, 2020) highlight the variation in traits based on recent environmental conditions, supporting the need to study bleaching under seasonally realistic conditions to more accurately predict an organism’s stress response (e.g. conducting bleaching studies at the timing of the peak of temperature for that region). Thus, high-frequency sampling of coral multivariate phenotypes through time and following bleaching stress is necessary to understand the mechanisms and sequential timing of the symbiotic state during stress and subsequent recovery.

Here, we tested the effect of an environmentally realistic extended heatwave in the context of diurnally fluctuating temperature and ocean acidification on coral physiology to improve our understanding of coral stress biology and forecasting reef futures. Specifically,we conducted a 4-month multi-stressor time-series experiment during the fall warming season in Kāne‘ohe Bay, O’ahu, Hawai’i, to assess the temporal dynamics during exposure to increased temperature and partial pressure of CO_2_ (pCO_2_) in addition to seasonal change, to compare differential stress response in a resilient and a sensitive coral species, and to investigate temporal dynamics during recovery and the potential for physiological legacies after an environmental perturbation.

## Methods

### Coral Collections

Coral fragments (one ∼3×3 cm fragment per colony from 75 colonies of *Montipora capitata* Dana 1846 and 75 colonies of Pocillopora acuta Lamarck 1816) were collected via clippers at ∼3–4.5 m depth from each of six reef areas ranging across the north to south span of Kāne‘ohe Bay including both fringing and patch reefs [Lilipuna Fringe: 21°25’45.9″N 157°47’28.0″W; Hawai‘i Institute of Marine Biology (HIMB): 21°26′09.8″N 157°47′12.7″W; Reef.11.3: 21°27′02.9″N 157°47′41.8″W; Reef.18: 21°27′02.9″N 157°48′40.1″W; Reef.35.36: 21°28′26.0″N 157°50′01.2″W; Reef.42.43: 21°28′37.9″ N 157°49′36.8″ W] under Hawai‘i Department of Aquatic Resources Special Activity Permit 2019-60, between 4 and 10 September 2018 (Fig. 1A,B). This collection approach, ensuring each fragment was >10 m apart from each other, was designed to target a diverse collection of genotypes that represent the coral population for *M. capitata* and *P. acuta* in Kāne‘ohe Bay. Coral fragments were brought back to HIMB and affixed in an upright position to individually numbered plastic coral fragment mounting plugs by hot gluing the base of the skeleton to the plug (McMaster-Carr, Hot-Melt Glue #7518A54). Coral fragments were allowed to acclimate for ∼14 days in outdoor mesocosm tanks and then were randomly allocated in an even distribution (n=405 *P. acuta* and n=440 *M. capitata*) to 12 replicate tanks for 2 months (weeks 1–8) of temperature and pCO_2_ stress exposure and 2 months of ambient condition recovery (weeks 9–16) as described below. The goal of our coral collection design was to examine the population of corals in Kāne‘ohe Bay. Therefore, the coral fragments were randomly allocated to the tanks, so the site of collection was not included statistically.

**Fig. 1.**
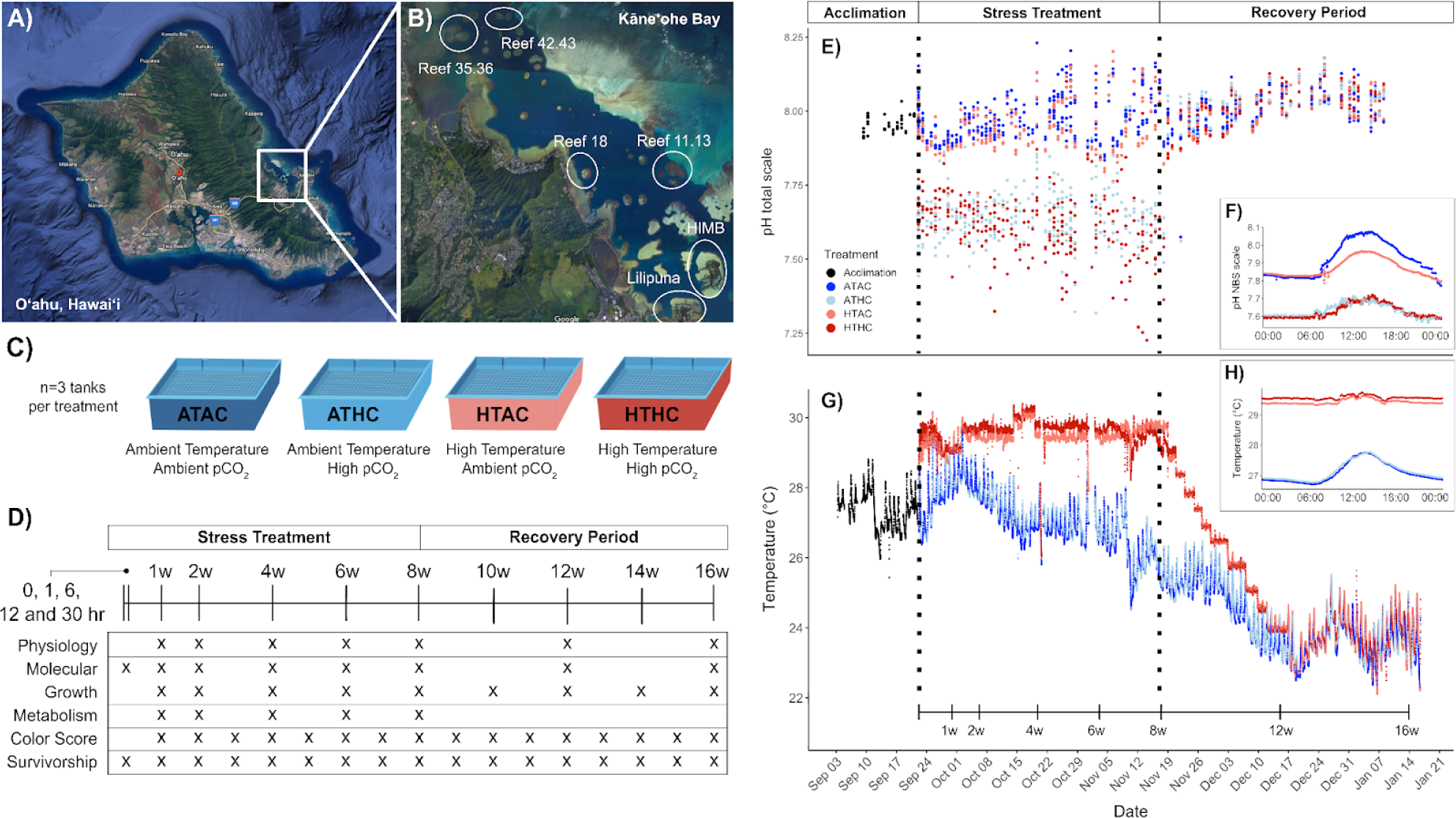
Experimental design for an environmentally and seasonally realistic extended heatwave scenario. (A) A map of O’ahu, Hawai‘i, with a highlight box around the study site. (B) A map of Kāne‘ohe Bay with six circled collection sites. (C) Twelve tanks (n=3 per treatment) were assigned a treatment with the following color and acronym code: ATAC (dark blue), ATHC (light blue), HTAC (light red), HTHC (dark red). (D) Sampling timeline with an ‘x’ to indicate when sampling occurred for each response category (n=6 physiology; n=3 molecular per species per treatment). Physiology includes antioxidant capacity, soluble protein, ash-free dry weight (AFDW), chlorophyll a (*chl a*) content and endosymbiont density. Metabolism includes photosynthetic and respiration rates. (E) Discrete measurements for pH treatment conditions (pH total scale) for the full experimental period. (F) Average daily pH treatment conditions modeled after natural fluctuations that occur in Kāne‘ohe Bay. Data come from continuous measurements from an Apex System set up during the first 2 weeks of the exposure period. (G) Discrete measurements for temperature treatment conditions (°C) for the full experimental period. A timeline of physiological sampling dates is indicated above the x-ais. (H) Average daily temperature treatment conditions modeled after natural fluctuations that occur in Kāne‘ohe Bay.

### Experimental Timeline

Coral fragments were exposed to one of four experimental conditions – (1) ambient temperature+ambient pCO_2_ (ATAC), (2) ambient temperature+high pCO_2_ (ATHC), (3) high temperature+ambient pCO_2_ (HTAC) and (4) high temperature+high pCO_2_ (HTHC) – for 2 months (22 September–17 November 2018; Fig. 1C). Treatment conditions (n=3 replicate tanks per treatment) were randomly assigned to 12 outdoor flow-through mesocosm tanks (122×122×30 cm; 510 l) supplied with sand-filtered seawater from Kāne‘ohe Bay. Flow rates, measured daily with a graduated cylinder and timer, averaged 84.36±1.20 ml s^−1^ (n=826), providing full mesocosm tank turnover every ∼2 h. Mesocosm tanks were 60% shaded from full irradiance, and photosynthetically active radiation (PAR) was measured continuously with an Apex cosine corrected PAR Sensor (PMK, Neptune Systems, accuracy =±5%) that was cross-calibrated to the Li-Cor cosine corrected PAR sensor (LI-193 spherical underwater quantum sensor, LI-COR Biosciences). Additionally, light values (roughly 100–700 PAR) from six different positions in each tank were compared to determine spatial differences within each tank. There was no significant difference in light between positions in each tank (n=4 per position per tank; position P=0.978; tank P=0.143). Based on these results, light was measured in the center of the tank approximately daily for the duration of the experiment using the LI-193 spherical underwater quantum sensor. To further reduce any potential position effects with respect to incoming water, heater position or bubble stream, the positions of the coral fragments in the tank were changed weekly. During this process, coral fragments were gently cleaned with toothbrushes and tanks were scrubbed to remove excess algal growth.

### Experimental temperature conditions

Temperature treatment conditions were programmed to mimic the natural daily fluctuations of the environment at the collection sites in Kāne‘ohe Bay, Hawai‘i (NOAA Moku o Lo‘e Buoy data from September 2018, https://www.ndbc.noaa.gov/station_page.php? station=mokh1; Fig. 1G; Fig. S1). Based on these data, the high temperature treatment fluctuated between ∼29 and 30°C to reflect previous marine heatwaves in Kāne‘ohe Bay, Hawai‘i (+2°C above ambient temperature). Temperature was monitored with Apex Extended Life Temperature Probes (accuracy =±0.05°C, Neptune Systems) and temperature loggers (HOBO Water Temp Pro v2, accuracy=±0.21°C, resolution=0.02°C, Onset Computer Corp.) that were placed in each tank at the same height as the coral fragments for the duration of the experiment and logged temperature at 10 min intervals. Temperature was separately controlled by submersible heaters (ProHeat D-1500 Heater Controllers, precision ±1°C). Ambient temperature treatments reflected the natural conditions of Kāne‘ohe Bay owing to the flow-through nature of the tanks.

### Experimental pCO_2_ conditions

Experimental pCO_2_ treatment conditions were based on measurements of ambient conditions in Kāne‘ohe Bay, Hawai‘i (Drupp et al., 2011), and projected future lower pH conditions in embayments, which can have lower mean pH and higher diel fluctuations owing to calcification, photosynthesis and respiration of the benthic community and increased residence time in the embayment (Jury et al., 2013; Shaw et al., 2016). The target pH range was based on values reported from low pH measured on the HIMB patch reef in Kāne‘ohe Bay (Guadayol et al., 2014). The daily fluctuating pH levels in the high pCO_2_ treatment tanks were maintained between 7.6 and 7.7 (pH NBS scale) with an independent pH-stat feedback system in each tank, and ambient conditions fluctuated between 7.9 and 8.0 (pH NBS scale; Fig. 1E,F). To generate the high pCO_2_ treatment, two 99.99% food-grade CO_2_ cylinders were connected to an automatic gas cylinder changeover system (Assurance Valve Systems, Automatic Gas Changeover Eliminator Valves #6091) to prevent an abrupt shortage of CO_2_ supply. CO_2_ was added into the seawater on-demand through gas flow solenoids (Milwaukee MA955), based on the pH reading (NBS) of a lab-grade pH probe (Apex pH, Neptune Systems) in each tank. CO_2_ was delivered via airlines plumbed into a Venturi injector (Forfuture-go G1/2 Garden Irrigation Device Venturi Fertilizer Injector), which was connected to a water circulating pump (Pondmaster Pond-mag Magnetic Drive Water Pump Model 5). Gas injected into the system was either CO_2_ or ambient air depending on treatment, and was injected through a Venturi injector connected to a pressure-driven pump, resulting in constant bubbling and moving water. An Apex AquaController (Neptune Systems) environmental control system with a wifi base unit (Apex Controller Base Unit, Neptune Systems) was linked to 12 individual monitoring units (Apex PM1 pH/ORP Probe Module, Neptune Systems) with Apex lab-grade pH Probes (accuracy=±0.01 pH NBS scale, Neptune Systems), which were used for microprocessor-control of a power strip (Apex Energy Bar 832, Neptune Systems) containing 12 individual solenoids. This pH-stat feedback system constantly monitored seawater temperature and pH conditions. Once sampling and treatments were completed, the pCO_2_ injections were shut off immediately in all tanks and the temperature gradually decreased over the course of a week (Fig. 1G), and fragments were exposed to ambient temperature and ambient pCO_2_ conditions for∼2months (18November 2018–12 January 2019), identified here as the recovery period.

### Total Alkalinity and Carbonate Chemistry

Tank parameters [temperature (°C), pH (total scale) and salinity (psu)] were measured 2–3 times daily using a handheld digital thermometer (Fisherbrand, Traceable Platinum Ultra-Accurate Digital Thermometer, accuracy=±0.05°C, resolution=0.001°C) and a portable multiparameter meter (Thermo Fisher Scientific, Orion Star A series A325). A pH probe (Mettler Toledo, InLab Expert Pro pH probe #51343101; accuracy=±0.2 mV, resolution=0.1 mV) and conductivity probe (Orion, DuraProbe 4-Electrode Conductivity Cell Model 013010MD; accuracy=0.5% of psu reading, resolution=0.01 psu) were used with an Orion A Star meter to measure pH (mV) and salinity (psu), respectively. pH (total scale) was calculated from standard curves of pH (mV) across a range of temperature (°C) in a tris standard (Dickson Laboratory, Tris batch T27 bottles 269 and 236, and batch T26 bottle 198). Water samples (125 ml) were taken from each tank twice a week to measure total alkalinity and, along with the pH on the total scale, as described above, to calculate carbonate chemistry. An automated titrator (Mettler Toledo T50) was used to titrate water samples with salinity-adjusted 0.1 mol l−1 hydrochloric acid (Dickson Laboratory Titrant A3, A14). A non-linear, least-squares procedure of the Gran approach (SOP 3b; Dickson et al., 2007) was used to calculate total alkalinity (TA; μmol kg^−1^ seawater). Accuracy of the titrations was determined using certified reference material [Dickson Laboratory CO_2_ Certified Reference Material (CRM) batch 132 (salinity=33.441, TA=2229.24±1.03 μmol kg^−1^), batch 137 (salinity=33.607, TA=2231.59±0.62 μmol kg^−1^), batch 176 (salinity=33.532, TA=2226.38±0.53 μmol kg−1); NOAA Ocean Carbon and Acidification Data System]. Tank total alkalinity values were only accepted if the total alkalinity value of the CRM from that day was within 1%of the reported CRM. Carbonate values, including aragonite saturation (Ω), carbon dioxide (CO_2_), carbonate (CO ^−3^), dissolved inorganic carbon (DIC), bicarbonate (HCO^3−^), partial pressure of carbon dioxide (pCO_2_) and pH, were calculated using SEACARB (v3.2.16 in R Studio; https://CRAN.R-project.org/package=seacarb) with the following parameters: calculated total pH, calculated TA, flag=8 [pH and ALK given], P=0, Pt=0, Sit=0, pHscale=‘T’, kf=‘pf’, k1k2=‘l’ and ks=‘d’. The equilibrium constant of hydrogen fluoride (mol kg−1) (kf=‘pf’) was set to ‘pf’ for salinities ranging from 10 to 40 psu and temperature ranging from 9 to 33°C (Perez and Fraga, 1987) and recommended by Dickson et al. (2007). The first (K1) and second (K2) dissociation constants of carbonic acid (mol kg−1) were set to ‘l’ (k1k2=‘l’) for salinities ranging from 19 to 43 psu and temperature ranging from 2 to 35°C (k1k2=‘l’; Lueker et al., 2000). The stability constant of hydrogen sulfate (mol kg−1) (Ks) was set to ‘d’ (Dickson, 1990) for salinities ranging from 5 to 45 psu and temperature ranging from 0 to 45°C. Salinity and temperature inputs were the measurements taken at the time of the water sample collection.

### Physiological Response Variables

Physiological sampling occurred at the following time points: 1, 2, 4, 6, 8, 12 and 16 weeks (Fig. 1D). Physiological samples were immediately snap-frozen in liquid nitrogen after photosynthetic and respiration rates were measured (described below; n=2 per tank×3 tanks per treatment=6 fragments per treatment per species per time point). All samples were stored at −80°C until subsequent processing.

### Survivorship, growth rates, and color score

Survivorship was tracked daily throughout the 2-month exposure and 2-month recovery periods using a binary score for dead versus alive. Alive was defined as any fragment with live tissue. Growth over time was determined by the buoyant weight technique (Davies, 1989; Jokiel et al., 1978) using air, freshwater and saltwater standard curves as a function of temperature.

Each coral fragment was buoyant weighed in its respective treatment temperature and pCO_2_ conditions. Wet buoyant mass was converted to dry mass using the aragonite density of 2.03 g cm^−3^ for Montipora spp. (Anthony and Hoegh-Guldberg, 2003) and 2.93 g cm^−3^ for *P. acuta* (Jokiel et al., 1978). Rates were normalized to surface area, determined by a single-dip wax dipping technique (Veal et al., 2010), and reported in units of mg CaCO_3_ cm^−2^ day^−1^. To assess tissue color change over time as a proxy for cell density and pigmentation sensu Edmunds et al. (2003), each coral fragment was photographed once a week (16 time points across the 4-month experiment) with a red, green, blue color standard. ImageJ was used to extract the mean red, green and blue color score for each coral fragment and these values were normalized to the mean of the red, green and blue color standards, respectively. The color scorewas quantified as principal component 1 (PC1; using the princomp function in R) from the principal component analysis (PCA) (sensu Edmunds et al., 2003).

### Photosynthesis and Respiration Rates

Prior to experimental exposures, 10 coral fragments (5 per species) were used to generate a photosynthetic irradiance (PI) curve to determine saturating irradiance for assessing rates of photosynthesis. Fragments were exposed to 10 light levels: 0, 15, 30, 60, 91, 136, 227, 416, 529 and 756 μmol photons m^−2^ s^−1^ (as measured by the Apex PAR sensor described above) generated by two LEDs (Aqua Illumination Hydra FiftyTwo) hung above the incubation chambers (described below). Rates of oxygen consumption or evolution were extracted using local linear regressions in the LoLinR package (Olito et al., 2017) in μmol O_2_ l^−1^ s^−1^. Rates were corrected to chamber volume including coral displacement. Rates of the blank chambers were subtracted from the rates of the coral fragments within each run to account for background photosynthesis or respiration owing to phytoplankton, zooplankton and bacteria. Rates of oxygen flux were normalized to surface area for final units of μmolO_2_ cm^−2^ h^−1^. Curve fitting of a non-linear least squares fit for a non-rectangular hyperbola (Marshall and Biscoe, 1980; Heberling and Fridley, 2013) was used to identify PI curve characteristics of each species. To calculate net photosynthesis (Pnet; μmol O_2_ h^−1^), the following PI curve fit equation was used:

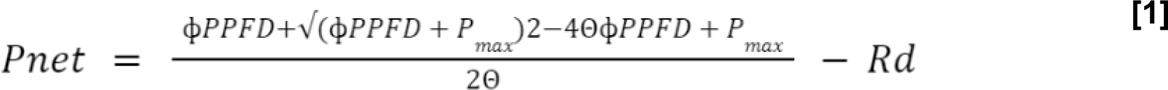

where PPFD (photosynthetic photon flux density) is the irradiance (μmol photons m^−2^ s^−1^), areal Pnet is net photosynthesis normalized to tissue area covered (cm^2^) and cellular Pnet is net photosynthesis normalized to symbiont cell density (number of cells), Pmax is the maximum gross photosynthetic rate (μmol O2 cm^−2^ h^−1^), φ is the apparent quantum yield (mol O_2_ mol photons^−1^) and Rd is the light-enhanced dark respiration rate (Edmunds and Davies, 1988) (μmol O2 cm^−2^ h^−1^).

Photoinhibition, as represented by Ik,was calculated as Pmax divided by the apparent quantum yield. PI curves were performed four times throughout the experiment to ensure the coral fragments were not reaching photoinhibition (Fig. S4). Fragments were placed in individual respiration chambers (∼610 ml), with individual temperature (Pt1000 temperature sensor, PreSens Precision Sensing GmbH) and fiber-optic oxygen probes (Oxygen Dipping Probes DPPSt7, accuracy=±0.05% O_2_, PreSens Precision Sensing GmbH) connected to a 10-channel oxygen meter (OXY-10 ST, accuracy= ±1.0°C, resolution=0.1°C, PreSens Precision Sensing GmbH) to evaluate photosynthesis (under saturating light conditions, as determined by PI curves described above) and light-enhanced dark respiration (Edmunds and Davies, 1988). The respirometry setup consisted of 10 chambers with stir bars and respective treatment water.

Samples were measured in a series of runs that consisted of eight fragments (n=4 per species) and two blank chambers per run exposed to PAR irradiance of ∼500 μmol photons m^−2^ s^−1^ for 15 min to assess photosynthetic rates. Immediately afterwards, these fragments were exposed to dark conditions (0 μmol photons m^−2^ s^−1^) for 20min to assess light-enhanced dark respiration. Following respirometry, fragments were immediately snap-frozen in liquid nitrogen and stored at −80°C. Raw oxygen flux data (temperature corrected) were thinned every 40 s and rates of oxygen flux were calculated with local linear regression as described above. Net photosynthesis was calculated by subtracting the oxygen change of the dark runs (a negative value) from the oxygen change of the light runs (a positive value), to isolate photosynthetic rates of the symbionts from the light enhanced dark respiration of the coral holobiont.

### Tissue removal, surface area, and endosymbiont density

Coral tissue was removed from the coral skeletons by airbrushing (Iwata Eclipse HP-BCS) with ice-cold 1× phosphate buffered saline (PBS) solution, and the total volumewas recorded. The tissue slurry was then homogenized for 15 s at high speed (PRO Scientific Bio-Gen PRO200 Homogenizer, 10 mm Homogenizer Probe). Of the total tissue slurry, aliquots of 500 μl were taken for endosymbiont density, 500 μl for host soluble protein and host total antioxidant capacity (TAC), 1 ml for chlorophyll a (*chl a*) concentration assays, and 5 ml for ash free dry weight (AFDW). The skeleton was rinsed in freshwater and soaked in 20% bleach for 24 h and dried at room temperature for 24 h. Symbiodiniaceae cell concentration was quantified with six technical separately loaded replicate counts on a hemocytometer (Hausser Scientific, Bright-Line Counting Chamber) after homogenizing the tissue slurry for 1 min. The host tissue was homogenized so that the symbiont cells were free of host tissue and clearly visible on the counting slide. The average count (cells ml^−1^) was standardized to the airbrushed homogenate volume and surface area to produce cell density values in cells cm^−2^.

### Host soluble protein and total antioxidant capacity

To quantify host soluble protein and host TAC, the host tissue was isolated from symbionts by centrifugation at 10,000 g for 10 min at 4°C. Soluble host protein of the supernatant was measured using the Pierce BCA Protein Assay Kit (Pierce Biotechnology, Waltham, MA, USA, catalog no. 23225) with two duplicates per sample measured by a 96-well microplate reader (BioTek Synergy HTX Multi-Mode Reader Model S1LFA) at a wavelength of 562 nm. Resulting values were calculated from a bovine serum albumin standard curve and standardized to homogenate volume and surface area to obtain host soluble protein values (mg cm^−2^). Host TAC was quantified in duplicate per sample with the Cell BioLabs OxiSelect Total Antioxidant Capacity (TAC) Assay Kit (catalog no. STA-360) according to the manufacturer’s instructions and compared with a uric acid standard curve. Antioxidants in the sample reduce Copper II to Copper I, resulting in positive correlation between copper reducing equivalents (CRE, mmol l^−1^) and antioxidant levels. CRE values were standardized to total soluble protein for final values in units of CRE μmol mg^−1^.

### Chlorophyll-a concentration

To isolate symbiont cells for *chl a* concentration measurements, a 1 ml aliquot of tissue homogenate was centrifuged at 13,000 g for 3 min and the supernatant was discarded. Then, 1 ml of 100% acetone was added to the symbiont pellet, mixed for 15 s on a lab vortex mixer, and incubated in the dark at 4°C for 24 h. Samples and acetone blanks were read in duplicate (200 μl each) on a 96-well microplate reader at λ=630 and 663 nm. *Chl a* concentrations were calculated with the following dinoflagellate equation for 100% acetone: *chl a*=11.43E663– 0.64E630 (Jeffrey and Humphrey, 1975). Concentration values were multiplied by 0.584 to correct for the path length of the plate for units of μg ml^−1^ and standardized to the airbrushed homogenate volume, normalized by surface area (μg cm^−2^) and per symbiont cell (μg cell^−1^).

### Tissue Biomass

Tissue biomass was quantified using the AFDW method. Specifically, 5 ml of tissue slurry was centrifuged at 13,000 g for 3 min. A volume of 4 ml of the supernatant host tissue was removed and placed in preburned aluminum pan weighed on an analytical balance (Mettler Toledo, model ML203T/00) and the symbiont pellet was resuspended in 1 ml PBS solution and added to separate pre-weighed burned aluminum pans. The samples were placed in a drying oven (Thermo Fisher Scientific, Heratherm General Protocol Oven, catalog no. 51028112) for 24 h at 60°C, weighed, and then placed in a muffle furnace for 4 h at 500°C (Thermo Fisher Scientific, Lindberg Blue M Muffle Furnace, catalog no. BF51728C-1). AFDW (mg cm^−2^) of the host and symbiont fractions was calculated as the post-drying oven mass (dry mass)–post-muffle furnace mass, and final values were standardized to the airbrushed homogenate volume and normalized to surface area. Symbiont to host (S:H) biomass ratios were calculated by dividing the symbiont AFDW by the host AFDW.

### Univariate statistical analysis

Owing to the near-complete mortality during the recovery period in *P. acuta*, only the exposure period was analyzed for the following physiological variables: tissue biomass, *chl a*, endosymbiont density, respiration rates, photosynthetic rates, host TAC and host soluble protein. *Montipora capitata* responses were analyzed in the exposure and recovery period separately to assess the effect of treatment on physiological state and the effect of treatment history on physiological state during a recovery period. Net photosynthetic rates per cm^2^ from the exposure period in *M. capitata* and color score for both species were square root transformed to meet assumptions of analysis. Gross photosynthetic rates per cm^2^ and growth rates in *M. capitata*, and growth rates and *chl a* per cell in *P. acuta* were log10 transformed for the exposure periods. Endosymbiont density, symbiont AFDW per cell, host soluble protein and host TAC were log transformed in both *M. capitata* and *P. acuta*. All physiological responses (tissue biomass, *chl a* concentration, endosymbiont density, respiration rates, photosynthetic rates, host TAC and host soluble protein) were analyzed with linear mixed-effects models (lme4 package version 1.1-27.1; Bates et al., 2015) with time, temperature and pCO_2_ as fixed factors, tank number as a random factor, and sum effect coding (Kugler et al., 2018) to test for interaction effects. The goal of our coral collection design was to examine the population of corals in Kāne‘ohe Bay. Therefore, the coral fragments were randomly allocated to the tanks, so site of collection was not included statistically, as there was not a balanced design across the tanks. Color score and growth rates were analyzed using linear mixed-effects models with time, temperature and pCO_2_ as fixed factors and plug ID nested within tank number as a random repeated measures factor. Survivorship probability was calculated with a Cox proportional hazards mixed effects model fit by maximum likelihood using the coxpH function in the coxme package (version 2.2-16; Therneau and Grambsch, 2013) with temperature and pCO_2_ treatments as fixed effects and plug ID as a random factor. Type III Wald χ2 ANOVA tests were performed on all models to assess significance of fixed effects. The normality and variance assumptions of the ANOVA test were assessed with quantile–quantile plots and Levene’s test. To determine baseline differences at week 1 in physiology at between the two species, treatment replicates were grouped for each response variable (n=6 per treatment×4 treatments=24 per species), prior to significant physiological change, and a Welch two-sample t-test was used for each univariate measurement to assess significance.

### Multivariate statistical analysis

A correlation matrix was calculated using the ‘corrplot’ function from the corrplot R package (https://CRAN.R-project.org/package=corrplot) to determine which univariate physiology variables were correlated and thus what to include in multivariate analyses. Gross and net photosynthetic rates were highly correlated and as a result, only net photosynthetic rates were used in downstream analyses as measure of symbiont photophysiology without the influence of cellular respiration. Multivariate physiological matrices were centered and scaled prior to principal components analyses that were performed using the ‘prcomp’ function from the stats package in R (https://www.r-project.org/) to examine multivariate physiological states for each species in the exposure period, as well as the recovery period for *M. capitata*. Permutational multivariate analysis of variance (PERMANOVA) tests were used to assess significant differences in multivariate physiology with time, temperature and pCO_2_ as fixed effects (‘adonis’ function in the R package vegan; https://CRAN.Rproject.org/package=vegan) separately for exposure and recovery periods for each species.

### Molecular response variables

Molecular sampling occurred at the following time points: 0, 6, 12 and 30 h, and 1, 2, 4, 6, 8, 12 and 16weeks (Fig. 1D). At each time point, one fragment was selected at random from each tank of each species (n=1 per tank×3 tanks per treatment=3 fragments per treatment per species) and placed in sterile whirlpak bags and snap-frozen in liquid nitrogen at 13:00 h each day (with the exception of 6, 12 and 30 h time points).

### DNA and RNA Extractions

DNA and RNA were extracted simultaneously from the coral fragments that had been snap-frozen and stored at −80°C. A small piece (∼0.5–1 cm) was clipped off using clippers sterilized in 10% bleach, deionized water, isopropanol and RNAse free water, and then placed in 2 ml microcentrifuge tube containing 0.5 mm glass beads (Fisher Scientific, catalog no. 15-340-152) with 1000 μl of DNA/RNA shield. A two-step extraction protocol was used to extract RNA and DNA for RNASeq and DNA methylation, respectively, with the first step as a ‘soft’ homogenization to reduce shearing of RNA or DNA. Tubes were vortexed at high speed for 1 and 2 min for *P. acuta* and *M. capitata* fragments, respectively. The supernatant was removed and designated as the ‘soft extraction’. In the second step, 500 μl of DNA/RNA shield was added to the bead tubes and placed in a Qiagen TissueLyser for 1 min at 20 Hz. The supernatant was removed and designated as the ‘hard extraction’. Subsequently, 300 μl of samples from both soft and hard homogenate were extracted individually with the Zymo Quick-DNA/RNA Miniprep Plus Kit (Zymo Research, catalog no. D7003T) following the protocol instructions, with the following modifications. For the DNA elution step, the first elution of 10 μl of warmed 10 mmol l^−1^ Tris HCl was added to DNA columns for a 15 min room temperature incubation and centrifuged at 16,000 g for 30 s. A second elution of 100 μl of warmed 10 mmol l^−1^ Tris HCl was added for a 5 min incubation and centrifuged at 16,000 g for 30 s. Broad-range DNA quantity (ng μl^−1^) was measured with a Thermo Fisher Qubit Fluorometer, and DNA quality was assessed using gel electrophoresis. Only the second elution of the ‘hard extraction’ DNA elution was used for downstream internal transcribed spacer 2 (ITS2) sequencing protocols and analysis.

### Internal Transcribed Spacer 2 (ITS2) Sequencing

Custom primers were ordered from Integrated DNA Technologies with ITS2 region locus-specific sequences based on Lajeunesse and Trench (2000) and Illumina adapters (in italics) to create our ITS2 forward primer (5′ GAATTG CAG AAC TCC GTG TCG TCG GCA GCGTCA GATGTG TATAAG AGACAG 3′) and ITS2 reverse primer (5′GGGATC CATATGCTTAAG TTCAGCGGGTGTCTCGTG GGC TCG GAG ATG TGT ATA AGA GAC AG 3′). PCR master mix solution included (per sample values): 16.6 μl of 2× Phusion HiFi Mastermix (Thermo Scientific, catalog no. F531s), 14 μl of DNase/RNase freewater, 0.6 μl of 10 μmol l^−1^ ITS2 forward primer, 0.6 μl of 10 μmol l^−1^ ITS2 reverse primer and 1 μl of 10 ng μl^−1^ DNA sample or 1 μl ofDNase/RNase freewater as a negative control. To avoid PCR amplification bias, each sample was run in triplicate on 96-well plates with 32 μl of master mix and 1 μl of 10 ng μl^−1^ DNA sample in each well. Samples were run on the following PCR settings: 3 min at 95°C, then 35 cycles of 30 s at 95°C, 30 s at 52°C and 30 s at 72°C, followed by 2 min at 72°C and held indefinitely at 4°C. The triplicate PCR products for each sample were pooled and stored at −20°C. The PCR products were run on a 2% agarose gel to confirm 360– 390 bp pair fragment length (330–360 bp ITS2 region and locus-specific primers+33 bp adapter length). Then, 45 μl of PCR product was sent to the RI-INBRE Molecular Informatics Core at the University of Rhode Island (RI, USA) for the index PCR and sequencing. There, PCR products were cleaned with NucleoMag beads (Takara Bio, catalog no. 744503) and then visualized using gel electrophoresis. A second round of PCR was performed with 50 ng DNA input for 8 cycles to attach Nextera XT indices and adapters using the Illumina Nextera XT Index Kit (Illumina, catalog no. FC-131-200×) and Phusion HF PCR master mix (Thermo Fisher Scientific, catalog no. F531L). Products were then cleaned with NucleoMag beads and visualized using gel electrophoresis (1.5% agarose gel, 110 V, 35 min). The quality of samples was assessed using an Agilent BioAnalyzer DNA1000 chip (Agilent, catalog no. 5067-1505) and quantification was performed using a Qubit fluorometer with a High Sensitivity Kit (Invitrogen, catalog no. Q32584) prior to pooling all samples. The final pooled library was quantified using qPCR (Roche LightCycler480) with the KAPA Biosystems Illumina Kit (KAPA Biosystems). Samples were analyzed using 2×250 bp paired-end sequencing on an Illumina MiSeq (Illumina).

### Bioinformatic analysis

ITS2 sequences fromrawFASTQ fileswere analyzed using SymPortal (symportal.org; Hume et al., 2019) on theUniversity of Rhode Island’s High Performance Cluster. This workflow used Miniconda3 v4.9.2 and SymPortal v0.3.21-foss-2020b to assign sequences as defined intragenomic variants (DIV) and ‘ITS2 Type Profiles’. A raw counts matrix of the ITS2 Type Profile output from SymPortal was exported and analyzed locally using R Software Version 4.0.3 in RStudio Version 1.4.1103. Relative abundance and a Bray–Curtis distance matrix were calculated using the ‘vegan’ package (Dixon, 2003). A PERMANOVA was performed for each species to assess significant effects of time point, treatment and their interaction.

## Results

At the end of the ∼14-day acclimation period, 89.8% of *P. acuta* (n=405) and 97.8% of *M. capitata* (n=440) fragments were alive. During the exposures, treatments showed no significant variation in TA across the 16-week experiment (treatment P=0.776; ATAC=2159.06 ±5.88, ATHC=2162.46±0.12, HTAC=2155.58±5.25, HTHC= 2156.79±0.058; Fig. S2). Additionally, light conditions were not significantly different between treatments (treatment P=0.558; ATAC=179.70±9.97, ATHC=186.54±10.20, HTAC=176.64±10.84, HTHC=165.12±10.41; Fig. S3). High temperature treatments during the 8-week stress period were maintained above 29°C (HTAC=29.47 ±0.004, HTHC=29.59±0.004; n=7691), whereas ambient conditions, dictated by flow-through water, steadily declined from 28°C to 26°C (Fig. 1G). Recovery temperature conditions reflected natural conditions of Kāne‘ohe Bay with declines from 26°C to 23°C by week 16 in mid-January (Fig. 1G,H).

High pCO_2_ treatments (ATHC, HTHC) were maintained between 7.6 and 7.7 pH total scale (ATAC=8.03±0.011, ATHC=7.67±0.014, HTAC=8.01±0.011, HTHC=7.64±0.019). During the stress period, high pCO_2_ treatments were characterized by significant decreases in aragonite saturation (Ω; treatment×time P<0.0001), carbonate (CO_3_ ^2−^; treatment×time P<0.0001) and pH (treatment×time P<0.0001), as well as increases in carbon dioxide (CO_2_; treatment P<0.0001), dissolved inorganic carbon (DIC; treatment×time P<0.0001), bicarbonate (HCO^3−^; treatment×time P<0.0001) and pCO_2_ (treatment P<0.0001; Fig. 1E,F, Fig. S2). During recovery, all treatments reflected the natural conditions of Kāne‘ohe Bay, ∼8 pH total scale (Fig. 1E). Additional univariate figures and summary statistical tables can be found at Open Science Framework (OSF) doi:10.17605/OSF.IO/Q9HMC.

### High temperature drives bleaching responses and low P. acuta survivorship

In high temperature treatments (HTAC and HTHC), both species displayed significant color change relative to ambient, starting at 4 weeks in *P. acuta* and 6 weeks in *M. capitata*. High temperature and ambient pCO_2_ (HTAC) and high temperature high pCO_2_ (HTHC) treatments drove this decrease in color score in the exposure period in *M. capitata* (time×temperature×pCO_2_ P=0.0004) and *P. acuta* (time×temperature P<0.0001; time×temperature×pCO_2_ P<0.0001). Additionally, *P. acuta* color score decreased significantly through the autumn season (time P<0.0001) and in high pCO_2_ treatments over time (time×pCO_2_ P<0.0001; Fig. S6). Following this visual bleaching response in the exposure period, both high temperature (HTHC, HTAC) and pCO_2_ treatments (ATHC) elicited decreased survivorship probability in both *M. capitata* (temperature P <0.0001; pCO_2_ P=0.0007) and *P. acuta* (temperature×pCO_2_ P<0.0001; Fig. S5). In the recovery period, *M. capitata* fragments that experienced high temperature treatments (HTHC and HTAC) increased in color score to ambient values by week 16 (time×temperature×pCO_2_ P=0.0124; Fig. S6). The decreases in survivorship probability for *M. capitata* stopped once treatment conditions were removed. However, *P. acuta* fragments did not recover color, and survivorship probability continued to sharply decrease (Fig. S5).

### High temperature decreased metabolic rates in both species, but P. acuta displays higher baseline photosynthetic rates

There were significant decreases during the stress period (September to November) regardless of treatment in both species for respiration (*M. capitata* time P=0.0005; *P. acuta* time P<0.0001) and photosynthetic rates (*M. capitata* Pnet cm^−2^ time P=0.0060; *P. acuta* Pnet cm^−2^ time P<0.0001; *P. acuta* Pnet cell−1 time P <0.0001). However, *P. acuta* displayed significantly higher baseline (as defined by week 1) areal (Pnet cm^−2^ P=0.0002; Pgross cm^−2^ P=0.0013) and per cell photosynthetic rates (Pnet cell^−1^ P=0.0045), as well as higher photosynthesis:respiration ratios (P:R P=0.0023) than *M. capitata* (Fig. 2). During the exposure period, high temperature, rather than high pCO_2_, caused a decrease in respiration rates in *M. capitata* (temperature P=0.0203) and *P. acuta* (time×temperature P=0.0412). Similarly, high temperature drove significant decreases in net photosynthetic rates in *M. capitata* (time×temperature P=0.0079) and *P. acuta* (time×temperature P<0.0001).When net photosynthetic rates were normalized per symbiont cell, treatments did not affect *M. capitata* fragments (time×temperature×pCO_2_ P=0.1833), but *P. acuta* fragments in high temperature treatments displayed a significant decrease by week 6 (time×temperature P=0.0011). Concomitantly, by week 6 (out of 8 exposure weeks), *P. acuta* fragments also displayed a significant decrease in P:R in high temperature treatments (time×temperature P=0.0139) and overall changes in metabolic rates were more drastic in *P. acuta* fragments.

**Fig. 2.**
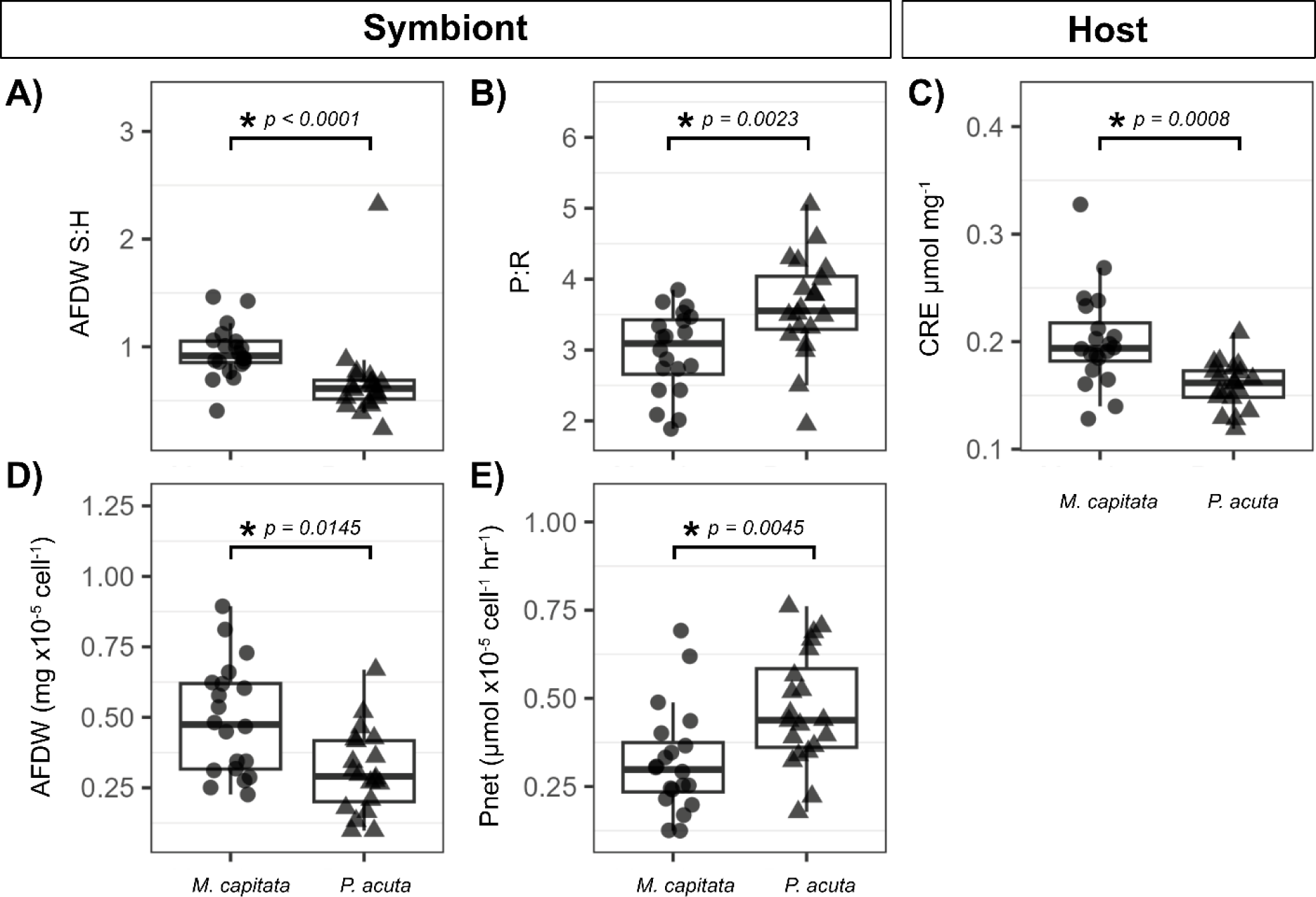
*Montipora capitata* and *Pocillopora acuta* baseline physiological measurements measured at week 1. (A) AFDW symbiont:host (S:H) ratio, (B) photosynthesis:respiration (P:R) ratio, (C) host total antioxidant capacity (CRE μmol mg−1), (D) symbiont cellular AFDW (×10−5 mg cell−1) and (E) cellular net photosynthetic rates (Pnet; ×10−5 μmol O2 cell−1 h−1).

### High pCO_2_ stress depresses seasonal increase in growth rates in M. capitata and high temperature decreases growth rates in both species

In both species, growth rates (mg CaCO_3_ cm^−2^ day^−1^) peaked in at week 4 of exposure (October) and gradually decreased by week 8 (November) of the exposure period (*M. capitata* time P<0.0001; *P. acuta* time P<0.0001) regardless of treatment. High temperature treatment decreased growth rates in later exposure time points (time×temperature×pCO_2_ P<0.0001) in both species. Similarly, both pCO_2_ and temperature treatments exhibited lower growth rates through the entire exposure period (time×temperature×pCO_2_ P<0.0001) in *P. acuta* fragments. In *M. capitata*, corals in high pCO_2_ treatments displayed significantly lower growth rates over time (Time× pCO_2_ P=0.0003). However, during the recovery period, *M. capitata* fragments that experienced only high pCO_2_ (ATHC) displayed increasing growth rates through week 16 (time× pCO_2_ P=0.0017), whereas fragments that experienced the high temperature treatments (HTAC and HTHC) gradually increased to ambient levels by week 16 (time×temperature P<0.0001).

### Host physiology: biomass remained stable while temperature stress increased TAC and decreased protein levels

On average, *M. capitata* exhibited higher baseline TAC than *P. acuta* (P=0.0008; Fig. 2). Host TAC (CRE μmol mg^−1^) remained constant in *M. capitata* through the exposure period regardless of treatment (time×temperature×pCO_2_ P=0.7028) but increased by week 8 in *P. acuta* under high temperature in HTAC and HTHC (time×temperature P=0.0277). During recovery, *M. capitata* fragments previously exposed to high temperature treatments showed reduced TAC at week 12, which returned to ambient treatment values by week 16 (time×temperature P=0.0237). Host soluble protein (mg cm^−2^) was not significantly different between species at baseline, week 1 (P=0.29), but significantly decreased through time from September to January in both *M. capitata* (time P<0.0001) and *P. acuta* (time P<0.0001). At week 6, protein concentration in *M. capitata* was significantly greater in both high pCO_2_ treatments (pCO_2_ P=0.0002) in comparison to HTAC and ATAC. Additionally, corals in high temperature treatments (HTHC and HTAC in comparison with ATAC and ATHC) displayed a decrease in protein over time (temperature×time P=0.0463). In *P. acuta* fragments, high temperature treatments decreased in protein at week 8 (time×temperature P=0.0490). Host AFDW (mg cm^−2^) was not significantly different at baseline, week 1 (P=0.22), or affected by treatment, but gradually and significantly decreased throughout the stress and exposure period from September to January in *M. capitata* (time P=0.0050) and *P. acuta* (time P=0.0188).

### Symbiont and holobiont physiology: stress treatments caused an increase in symbiont biomass but decreased chl a content and endosymbiont density

Symbiont biomass (mg AFDW cm^−2^) was unaffected by either temperature or pCO_2_ treatment (time×temperature×pCO_2_; *M. capitata* P=0.3839, *P. acuta* P=0.3790), but significantly decreased through the autumn season in both *M. capitata* (P=0.0013) and *P. acuta* (P=0.0028). Despite no treatment effect on symbiont and host biomass (mg AFDW cm^−2^), for *P. acuta* the symbiont:host biomass ratio significantly decreased in high temperature treatments (temperature P=0.0180) and decreased also over time (Time P=0.0090). *Montipora capitata* fragments exhibited significantly higher baseline week 1 biomass (mg cm^−2^ P=0.0145; mg cell^−1^ P=0.0079; Fig. 2) than *P. acuta*. In *M. capitata*, symbiont cellular biomass (mg AFDW cell^−1^) significantly increased in corals in the high temperature treatments during exposure (temperature×pCO_2_ P=0.0133) and then returned to ambient levels by week 16 (time×temperature P=0.0041). In *P. acuta*, symbiont per cell biomass decreased from September to January (P=0.0141), but significantly increased under both temperature and pCO_2_ exposure (time×temperature P<0.0001; time×pCO_2_ P=0.0146). Holobiont *chl a* (cm^−2^) increased over the autumn season in *M. capitata* (time P=0.0003; Fig. 3G), but high temperature exposure resulted in decreased *chl a* over time (time×temperature P=0.0089) that remained low in the recovery period (temperature P<0.0001). In *P. acuta*, *chl a* decreased over time (time P<0.0001) and in high temperature treatments over time (time×temperature P<0.0001; Fig. 3H). In *M. capitata*, only time significantly affected cellular *chl a* cell^−1^ (time P=0.0347), and in all fragments, regardless of treatment, *chl a* cell^−1^ increased from September to January. However, in *P. acuta*, *chl a* per cell was significantly affected by temperature and time (time×temperature P=0.0764). Endosymbiont density (cells cm^−2^) decreased in high temperature and pCO_2_ treatments over time in *M. capitata* (time×temperature P=0.0100; time×pCO_2_ P=0.0249) and the magnitude of decline was greater in high temperature in *P. acuta* (time×temperature P<0.0001). *Pocillopora acuta* endosymbiont density decreased by week 4 in all treatments (time P<0.0001; Fig. 3F), but all *M. capitata* fragments displayed an increase in densities in the same time period, followed decreased values by week 8 (time P<0.0001; Fig. 3E). By the end of the recovery period, *M. capitata* fragments that had experienced high temperature treatments did not fully recover in symbiont cell densities (temperature P=0.0070), despite being statistically indistinguishable in color score.

**Fig. 3.**
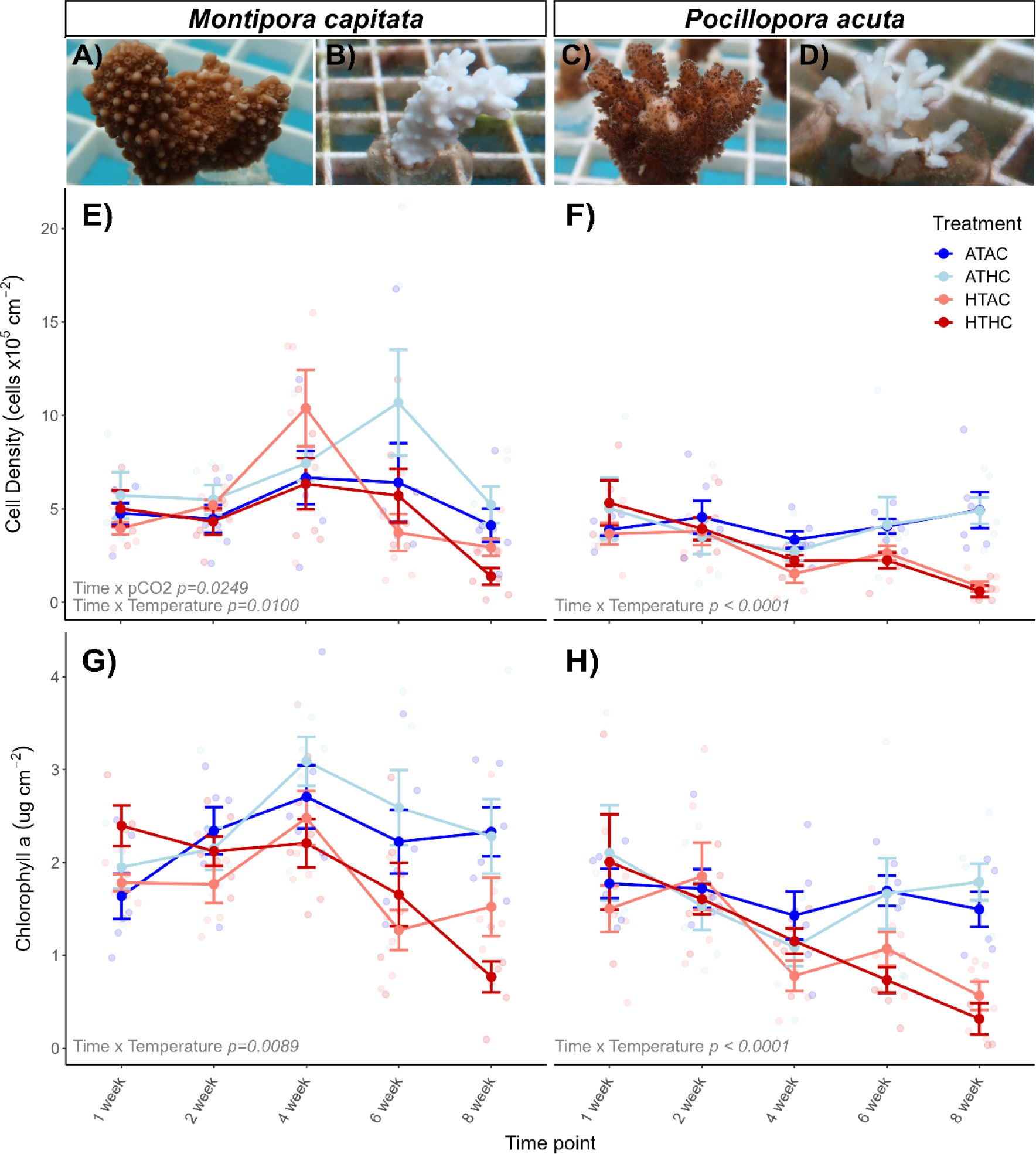
Holobiont chlorophyll a content (μg cm−2) and endosymbiont cell density (×105 cells cm−2) levels during the exposure period (weeks 1–8). The x-axis of E–H is time in weeks with sampling at 1, 2, 4, 6 and 8 weeks. Color indicates the four treatments. Points represent means±s.e.m., and significant effects from Type III ANOVA of linear mixed models for each response are labeled with corresponding P-value. (A–D) Examples of (A,C) healthy and (B,D) bleached *M. capitata* and *P. acuta* fragments. (E,F) Cell density levels (×105 cells cm−2) of (E) *M. capitata* and (F) *P. acuta*. (G) *Chl a* content (μg cm−2) of (G) *M. capitata* and (H) *P. acuta*.

### Multivariate physiology: high temperature exacerbates seasonal changes during stress and drives physiological states during recovery

In both species, treatments exacerbated existing seasonal shifts in multivariate physiological states (Fig. 4). In *P. acuta*, physiological state shifted over time (P<0.001) and in high temperature treatments (time×temperature P<0.001). Similarly, *M. capitata* fragments experienced a shift in physiological state over time (time P<0.001) owing to temperature (temperature P=0.005), and this was different between pCO_2_ treatments over time (time×pCO_2_ P=0.038). PC1 explained 39.83% of the variance and separated the samples between ambient temperature and high temperature for *P. acuta*, with changes in respiration rates and host TAC levels contributing most strongly to the separation (Fig. 4A,B). The PC2 axis explained 18.40% of the variation and the separation between treatment groups was characterized by changes in endosymbiont density and cellular symbiont biomass. In *M. capitata*, the PC1 axis explained 30.51% of the variation and was characterized by changes in the host (protein, biomass, respiration) and holobiont physiology (areal net photosynthetic rates, areal symbiont AFDW and holobiont *chl a* content) that contribute to separation primarily between time points. The PC2 axis explained 25.81% of the variation and was characterized by symbiont functions (symbiont AFDW mg cell^−1^, net photosynthetic rates and *chl a* content) that separate primarily treatment groups (Fig. 4C,D). During the recovery period, both time (P<0.001) and high temperature treatment (P<0.001) significantly affected physiological shifts in *M. capitata* (Fig. 5C,D). PC1 explained 37.61% of the variation and was characterized by host (biomass, protein) and holobiont traits (mg AFDW cm^−2^, *chl a* cm^−2^) that separate between the history of high temperature treatment groups in *M. capitata*. PC2 explained 30.11% of the variation and changes in endosymbiont density and symbiont traits (*chl a* cell^−1^, S:H biomass ratio, biomass) contributed strongly to the separation of treatment and time groups.

**Fig. 4.**
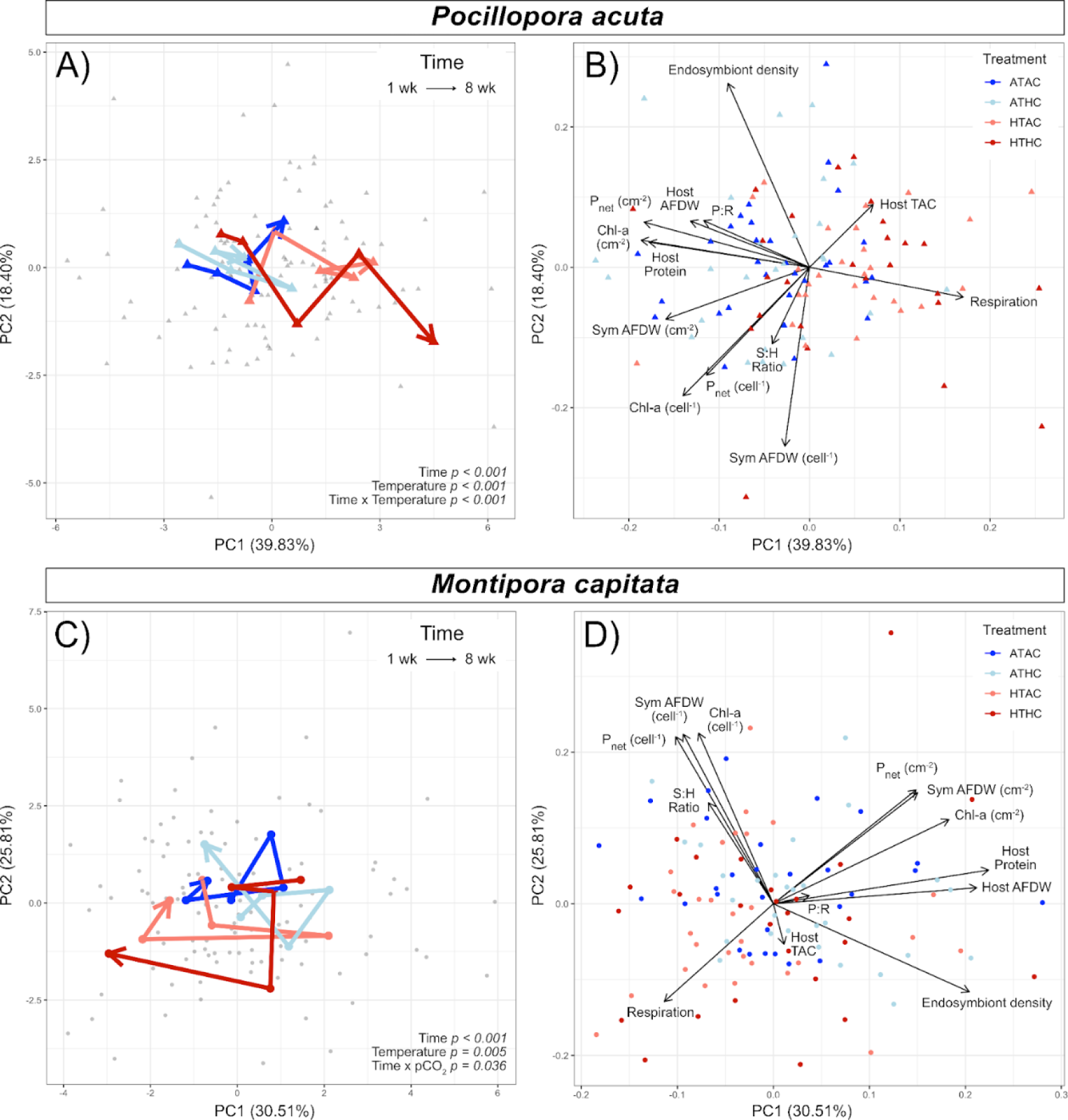
Principal component analyses of multivariate physiological state during the exposure period (weeks 1–8) for both species. (A,B) *Pocillopora acuta* fragments. (C,D) *Montipora capitata* fragments. Each point represents an individual coral fragment’s multivariate physiological state composed of 13 host and symbiont response variables and colored by treatment. Significant effects from PERMANOVA tests for each species are included within A and C (bottom right of each panel). PC axis labels indicate the percent variance explained by that axis. (A,C) Centroid multivariate state for each treatment at each time period is indicated by a bold, treatment-colored point. All mean points are connected within each treatment to show the shifts through time (week 1→8). (B,D) Biplots of the response variables are shown on the multivariate plots. Each black arrow represents a physiological response variable. The direction and length of each arrow indicates the relative strength of that variable to the PC axes.

**Fig. 5.**
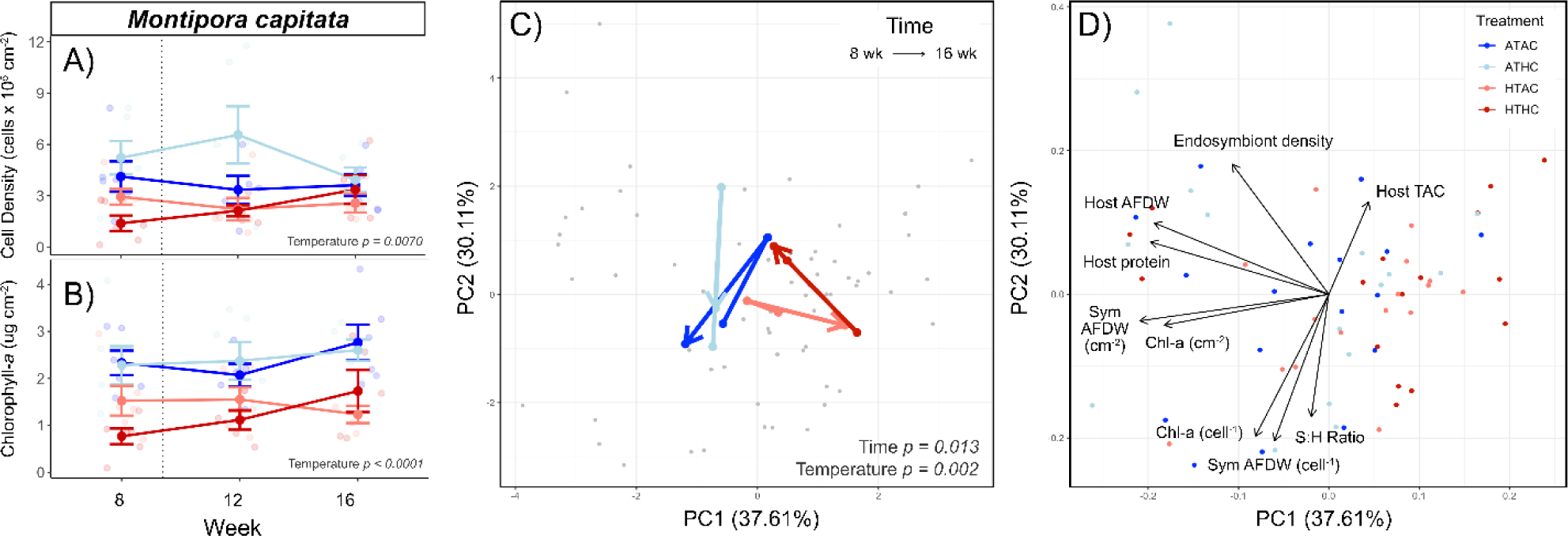
Bleaching assessment and principal components analysis for multivariate physiological state of *M. capitata* fragments during the recovery period (weeks 8–16). (A) Endosymbiont densities (×105 cells cm−2) and (B) Holobiont *chl a* content (μg cm−2) for the recovery period. (C,D) Each point represents an individual fragment’s multivariate physiological state composed of nine host and symbiont response variables (respiration and photosynthetic rates were not measured during the recovery period). (C) Centroid multivariate physiological state through time (week 8→16). (D) Biplot of individual physiological variable contributions to PCA separation. Each point (individual fragment) is colored by treatment. Significant effects from PERMANOVA tests for each species are indicated in gray text.

### Montipora capitata and P. acuta harbor distinctly different Symbiodiniaceae community compositions that do not change significantly during stress or recovery

The ITS2 sequencing generated 5,807,744 total raw reads from 251 samples, with an average of 22,775 reads per sample. The SymPortal analysis resulted in 4,902,971 total cleaned sequences, which excludes week 16 in *P. acuta* because of low sample sizes. *Montipora capitata* fragments harbored mixed Symbiodiniaceae communities composed of ITS2 symbiont type profiles with majority sequences ‘D1/D4’ and variations of ‘C17d/No-Name’ (Durusdinium and Cladocopium spp.; LaJeunesse et al., 2018; Fig. 6). *Pocillopora acuta* communities were dominated by type profiles in clade C with majority groups ‘C1d/C42.2’ and ‘C1d’. Endosymbiont communities were significantly different between species (PERMANOVA P<0.001) and differed only between two time points in *M. capitata* (12 h versus 24 h P=0.045) but did not significantly change in exposure treatments (P=1.000).

**Fig. 6.**
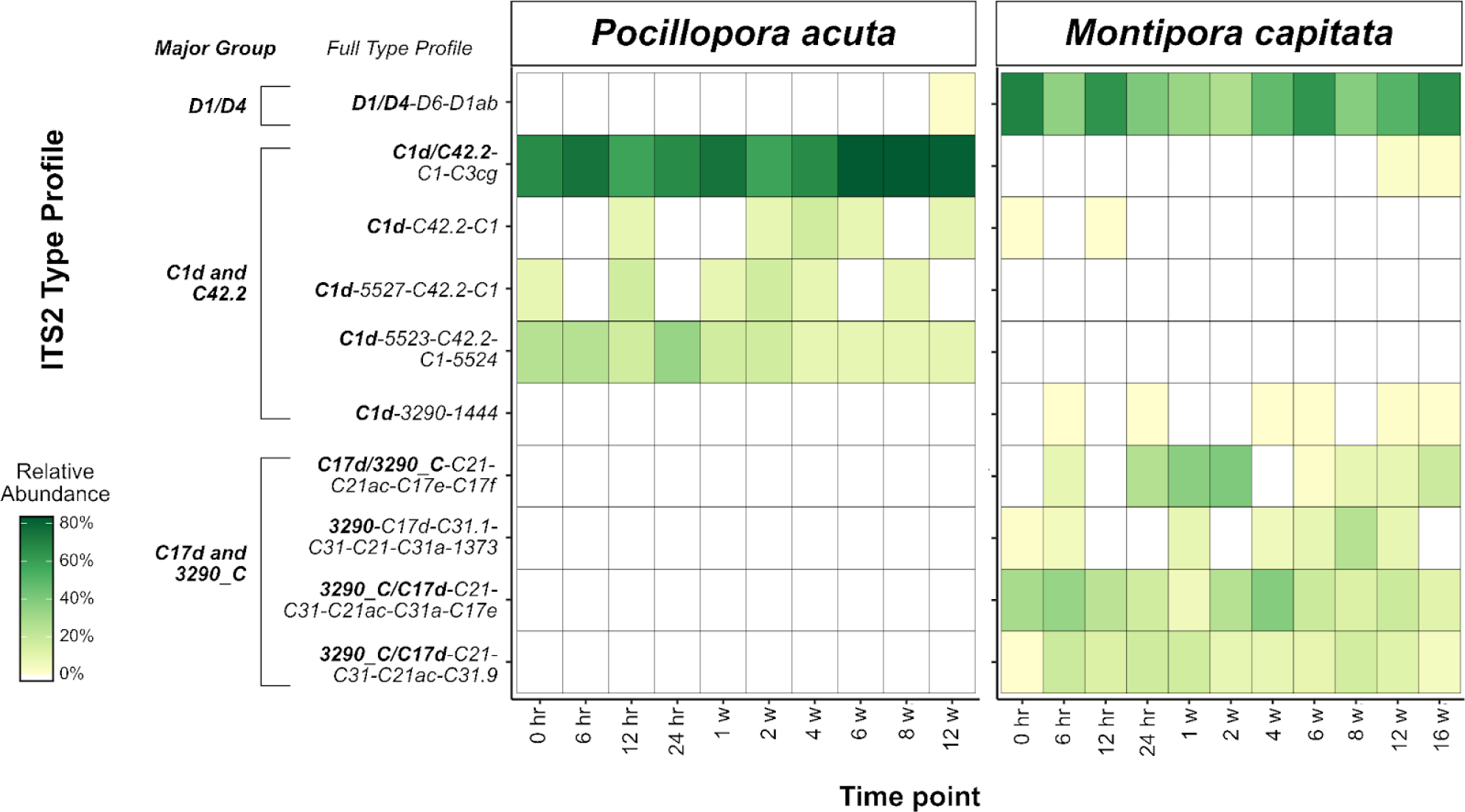
Relative abundance of Symbiodiniaceae ‘type profiles’ identified by ITS2 sequencing for both species during exposure and recovery periods. The y-axis includes each type profile, composed of a major sequence ID (in bold) and subsequent full type profile ID. Relative abundance is indicated by 0–100% continuous color scale, with the darkest green coloring indicating 100% abundance of that full type profile. The exposure period includes 0, 6 and 12 h, and 1, 2, 4, 6 and 8 weeks. The recovery period includes 12 and 16 weeks, but there was not a high enough sample size for *P. acuta* fragments at 16 weeks.

## Discussion

### Extended heatwave scenario induces species-specific mortality of corals and creates altered physiological legacies for the survivors

Scaling up from controlled single variable lab experiments to environmentally relevant multi-stressor experiments is essential to capture the effects of stressors in the context of seasonal physiological and thus the time-dependent and environmentally relevant sensitivity of corals to global change stressors. Our examination of the effects of high pCO_2_ and an extended heatwave at the peak of seasonal temperatures on *M. capitata* and *P. acuta* reinforce that increased temperatures are the more substantial threat to coral physiological performance (Putnam et al., 2013), survivorship (Klein et al., 2022) and physiological legacy effects (Wall et al., 2021) than high pCO_2_ conditions. We found that high pCO_2_ concentrations alone can elicit potentially beneficial effects in the more resilient *M. capitata* such as higher endosymbiont density and thus higher photosynthetic rates (cm^−2^) during exposure, and higher growth rates during recovery periods. However, these effects are not seen when high pCO_2_ conditions occur simultaneously with higher temperatures. The extent of physiological shifts, decreased survival probability and increased bleaching were all greater in *P. acuta* than in *M. capitata*, reinforcing the outcome of species-specific susceptibility and loss of corals on future reefs (Loya et al., 2001; van Woesik et al., 2011). Our data indicate that a combination of Symbiodiniaceae community and baseline levels of host antioxidant capacity and cellular photosynthetic rates are contributing to species-specific stress responses. Further, our findings of different physiology trajectories of *M. capitata* during the recovery period, specifically in those exposed to high temperature, support prior work that high temperature conditions leave lasting physiological legacies several months into recovery (Rodríguez-Troncoso et al., 2016; Schoepf et al., 2015; Rodrigues and Grottoli, 2007), and those legacies can have cumulative effects that shift physiological baselines leading into a subsequent stress event (Wall et al., 2021; Safaie et al., 2018; Ainsworth et al., 2016).

### Symbiodiniaeceae communities and high levels of antioxidants relative to cellular photosynthetic rates contribute to M. capitata thermal tolerance

Symbiodiniaceae communities aid in a coral holobiont’s tolerance and survival (Baskett, et al., 2009; Oliver and Palumbi, 2011); specifically, studies have documented that the presence of Durusdinium spp. (Silverstein et al., 2015; Guest et al., 2016) and/or mixed endosymbiont communities (Abbott et al., 2021) can be beneficial for coral holobiont survival. Our results support these hypotheses, as the mixture of Durusdinimum spp. (D1/D4; ∼60–80%) and Cladocopium spp. (C17/C31; ∼20–40%) in *M. capitata* was associated with higher thermal tolerance, compared with *P. acuta*’s Cladocopium spp. (C1; 100%) dominated communities. Although there was no evidence of Symbiodiniaceae community shuffling by treatment or through time, a holobiont’s community prior to an exposure event likely plays a crucial role in their ability to handle changing environmental conditions. Through these community composition differences, symbiont physiology influences holobiont tolerance (Hoadley et al., 2019; Ziegler et al., 2015), thus indicating that both the host and symbiont physiological states are critical to understanding the stress response of the holobiont (Bellantuono et al., 2012). Differential photophysiology, carbon assimilation and translocation to the host (Stat et al., 2008) and metabolite profiles (Matthews et al., 2017, 2018) have been seen between Symbiodiniaceae genera and thus influence holobiont tolerance (Wall et al., 2020; Matthews et al., 2018; Roach et al., 2021).

For both host and symbiont partners, a core physiological response to stress is the activity of antioxidant systems to neutralize ROS molecules that build up during daily metabolism and in response to stressful conditions such as increased temperature (Baird et al., 2009; Downs et al., 2000). These excess ROS molecules can leak into the host to cause a downstream cascade of cellular damage and eventually coral bleaching and apoptosis (Weis, 2008). A mismatch between the amount of symbiont-derived ROS molecules and the host’s antioxidant capacity may lead to higher susceptibility for coral bleaching (e.g. Cunning and Baker, 2013). We hypothesize that *P. acuta*’s naturally higher cellular photosynthetic rates, but lower host antioxidant capacity per unit protein, may lead to a faster build-up of ROS damage and consequently higher mortality rates. Thus, *P. acuta* are challenged under thermal stress because the symbiotic community is operating closer to the threshold between productivity and damage. In comparison, *M. capitata*’s naturally lower photosynthetic rates and higher host antioxidant capacity levels may be providing a greater buffer, allowing them capacity to support and balance their higher endosymbiont densities during November (Fig. 3), resulting in a more muted bleaching response when faced with stressful conditions. Further, *M. capitata* have the benefit of thicker tissues (Loya et al., 2001; Magnusson et al., 2007) with greater energy reserves (Rodrigues and Grottoli, 2007) in addition to their symbiont community composition. By the end of the exposure period, *P. acuta* host antioxidant capacity levels increased, and cellular photosynthetic rates decreased, but visual bleaching responses preceded this cellular physiological change, indicating that the response was insufficient. *Montipora capitata* may be well equipped to acclimatize to changing environmental conditions because of baseline holobiont physiological states and existing Symbiodiniaceae communities.

### Thermal exposure influences bleaching recovery trajectories in M. capitata

Recent studies have emphasized the importance of recovery time in between marine heatwaves (Osborne et al., 2017; Neal et al., 2017; Grottoli et al., 2014). Recovery time can be highly variable, with species taking anywhere from 1.5 months (Rodrigues and Grottoli, 2007), 2 months (Cunning et al., 2016), 3 months (Wall et al., 2019) up to 2 years (Matsuda et al., 2020). We found that in our experimental recovery period of 2 months following exposure, physiological metrics that decreased in chronic high temperature conditions stabilized as *M. capitata* fragments began to recover pigmentation (color score, symbiont density and *chl a*). Despite full recovery in color score and growth rates by week 16, *chl a* and endosymbiont densities did not fully recover to the same extent as the ambient condition corals, with differential physiological states and trajectories throughout the recovery period (Fig. 5). Our results indicate that extended high temperature exposure at the peak of seasonal temperature leaves a physiological legacy that lasts beyond 2 months. Even in this resistant and resilient species, legacy effects of bleaching on organismal state can be as long as 1–2 years (Wall et al., 2019, Roach et al., 2021; Thomas and Palumbi, 2017).

These legacy effect results are consistent with those of other studies that suggest that environmental conditions shift coral phenotypes, ultimately eliciting a lasting legacy that could have either a beneficial or detrimental priming effect for response in future stress events (Wall et al., 2021; Hackerott et al., 2021; Putnam et al., 2017; Brown et al., 2000). However, recovery time is imperative to understand whether the lasting effects of prior thermal events can be beneficial. Sublethal temperatures that induce a bleaching response 10 days prior to a larger intensity bleaching event have been shown to be beneficial (termed a ‘protective trajectory’; Ainsworth et al., 2016). Although beneficial effects were documented in long-term recovery periods of 11 months (Schoepf et al., 2015) following thermal perturbation, not all species recovery is on the same timeline, with Porites divaricata and Orbicella faveolata recovering faster than Porites astreoides (Schoepf et al., 2015). This potential for ‘environmental memory’ based on previous environmental conditions is clear in corals (Brown et al., 2000, 2022). Utilizing this to elicit purposeful stress hardening or environmental priming (Hackerott et al., 2021; Putnam et al., 2017; Brown et al., 2000; DeMerlis et al., 2022) is a proposed restoration technique (van Oppen et al., 2015), but it is clear there are limits of recovery time. *Montipora capitata* is likely to survive future thermal stress events, but back-to-back bleaching events and decreased recovery time may reduce their buffer and push their physiology closer to a threshold. More multi-year studies to track differential bleaching patterns over time are needed to elucidate the threshold between eliciting beneficial ‘environmental memory’ versus accumulated physiological damage and energetic deficits (Schoepf et al., 2015).

## Conclusion

This study highlighted the divergent thermal tolerance and recovery potential of *M. capitata* and *P. acuta*, indicating a potential for species-specific susceptibility to stress and loss of corals in future, more frequent and higher intensity bleaching events. *Montipora capitata* elevated stress tolerance may be due in part to mixed Cladocopium and Durusdinium Symbiodiniaceae communities, higher antioxidant capacity and elevated photosynthetic rates at baseline, and stability of metabolic and symbiont physiology during chronic high temperature conditions. In the 2 months of ambient recovery conditions that followed 2 months of elevated temperature and pCO_2_ conditions, *M. capitata* multivariate physiology did not fully recover. High temperature treatment history elicited a lasting multivariate physiological legacy that set recovering *M. capitata* on a different trajectory than those that experienced ambient conditions. This study illuminates and emphasizes the importance of environmentally and seasonally realistic time-series experiments to further understand the impacts of pre-existing seasonal dynamics on thermal stress response. Multi-year repeated bleaching event studies are needed to disentangle howa physiological legacy may be adaptive or maladaptive under predicted climate change scenarios.

## Contributions

Conceptualization: E.L.S., H.M.P.; Methodology: E.L.S., K.H.W., A.F., S.G., A.M., H.M.P.; Software: E.L.S., H.M.P.; Validation: E.L.S., H.M.P.; Formal analysis: E.L.S., H.M.P.; Investigation: E.L.S., H.M.P.; Resources: E.L.S., H.M.P.; Data curation: E.L.S., K.H.W., A.F., S.G., A.M., H.M.P.; Writing - original draft: E.L.S., H.M.P.; Writing - review & editing: E.L.S., K.H.W., A.F., S.G., A.M., H.M.P.; Visualization: E.L.S., H.M.P.; Supervision: E.L.S., H.M.P.; Project administration: E.L.S., H.M.P.; Funding acquisition: H.M.P.

## Data Availability

All data and code used to produce results and complete analyses are available at https://github.com/hputnam/Acclim_Dynamics. Laboratory protocols and processing can be found on Emma Strand’s Open Laboratory Notebook at https://emmastrand.github.io/EmmaStrand_Notebook/. ITS2 Sequencing FASTQ files are uploaded to NCBI under Project #PRJNA761780.

## Acknowledgements

We thank the Hawai‘i Institute of Marine Biology and the Coral Resilience Laboratory for hosting us to conduct the experiment on Coconut Island, and Chris Suchoki and Adam Helbig for assistance with field work. Ariana Huffmyer provided valuable feedback on an earlier version of the manuscript. As guests, we recognize and give thanks for the land and water resources of the ‘ā ina and the traditional owners of the land, kānaka ‘ō iwi, on which this experimental work was conducted in the Kāne‘ohe Ahupua’a and the islands of Hawai’i. The authors acknowledge use of the computational resources of the URI Center for Computational Research for this work.

---

**Fig. S1.**
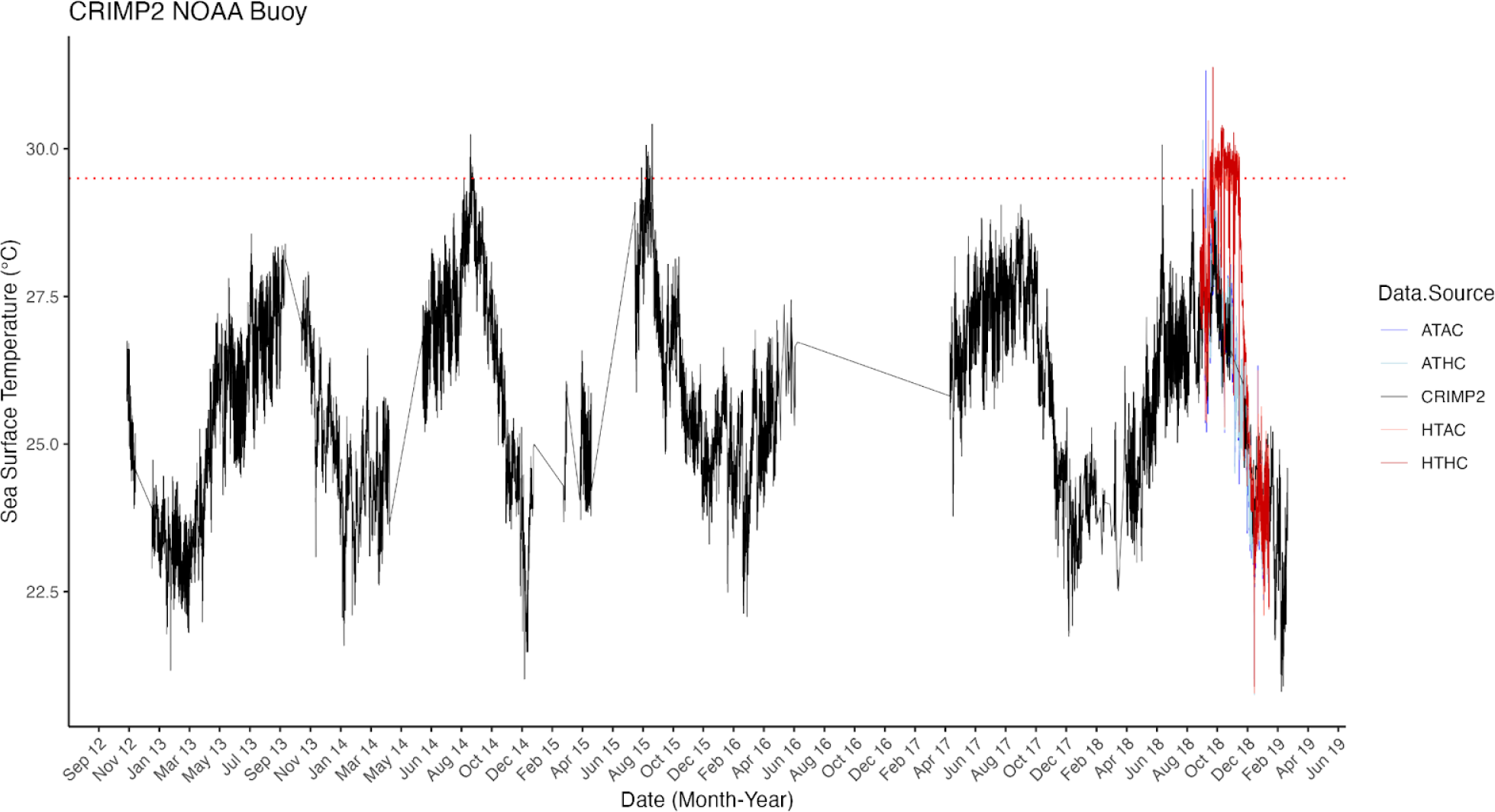
NOAA Moku o Loʻe Buoy data (CRIMP2) from September 2012 - June 2019 with treatment conditions overlayed. CRIMP2 buoy data (sea surface temperature °C) was used to determine an environmentally realistic range of temperatures for the experimental exposure period. The red dotted line represents the average high temperature treatment condition. CRIMP2 information can be found at https://www.ndbc.noaa.gov/station_page.php?station=mokh1.

**Fig. S2.**
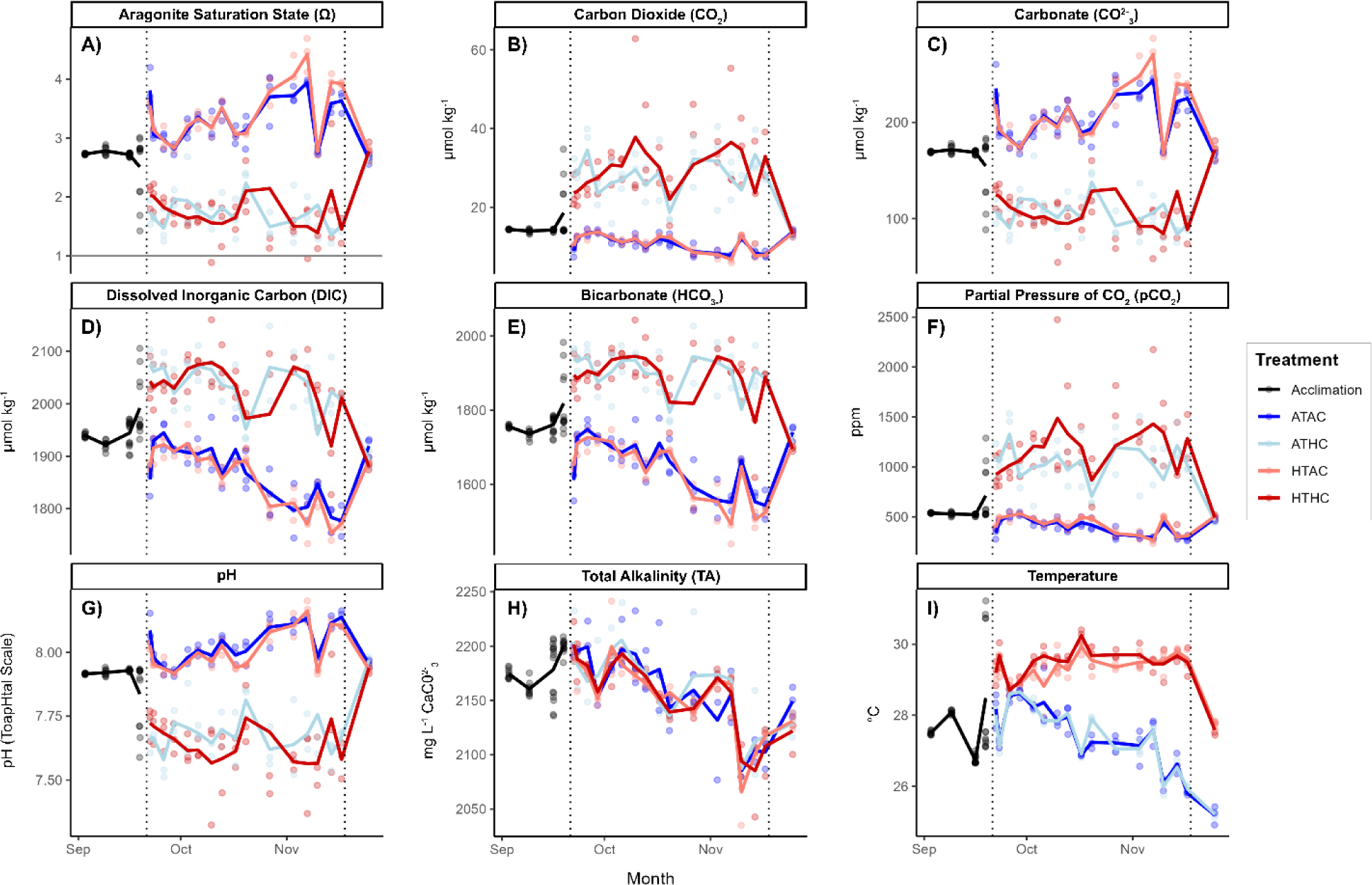
Carbonate chemistry parameters during the acclimation, exposure, and beginning of the recovery period for all treatments. Treatment is indicated by color and vertical dotted lines mark the start and end of the treatment period. All x-axes represent time by month. Parameters include the following: (A) Aragonite Saturation State (Ω). (B) Carbon dioxide (CO_2_; μmol kg^-1^). (C) Carbonate (CO^2-^_3_; μmol kg^-1^). (D) Dissolved inorganic carbon (DIC; μmol kg^-1^). (E) Bicarbonate (HCO_3-_; μmol kg^-1^). (F) Partial pressure of CO_2_ (pCO_2_; ppm). (G) pH (total scale). (H) Total alkalinity (TA; mg L^-1^ CaCO_3_) (I). Temperature (°C).

**Fig. S3.**
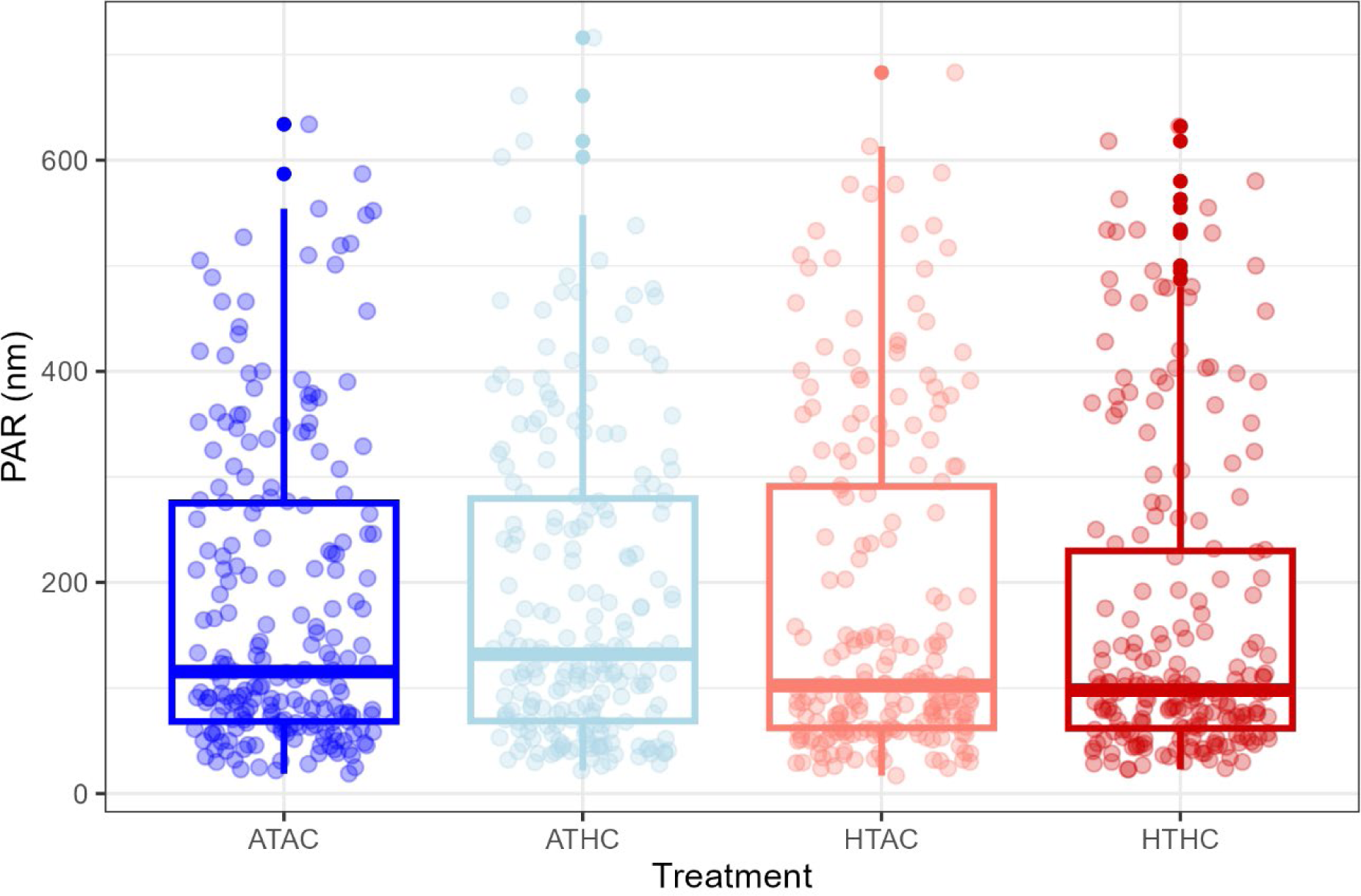
Light data from the exposure period for all treatments. Light reported in PAR (nm) measured with a LICOR-193 as described in methods. Color indicates treatment (dark blue = ATAC, light blue = ATHC, light red = HTAC, and dark red = HTHC).

**Fig. S4.**
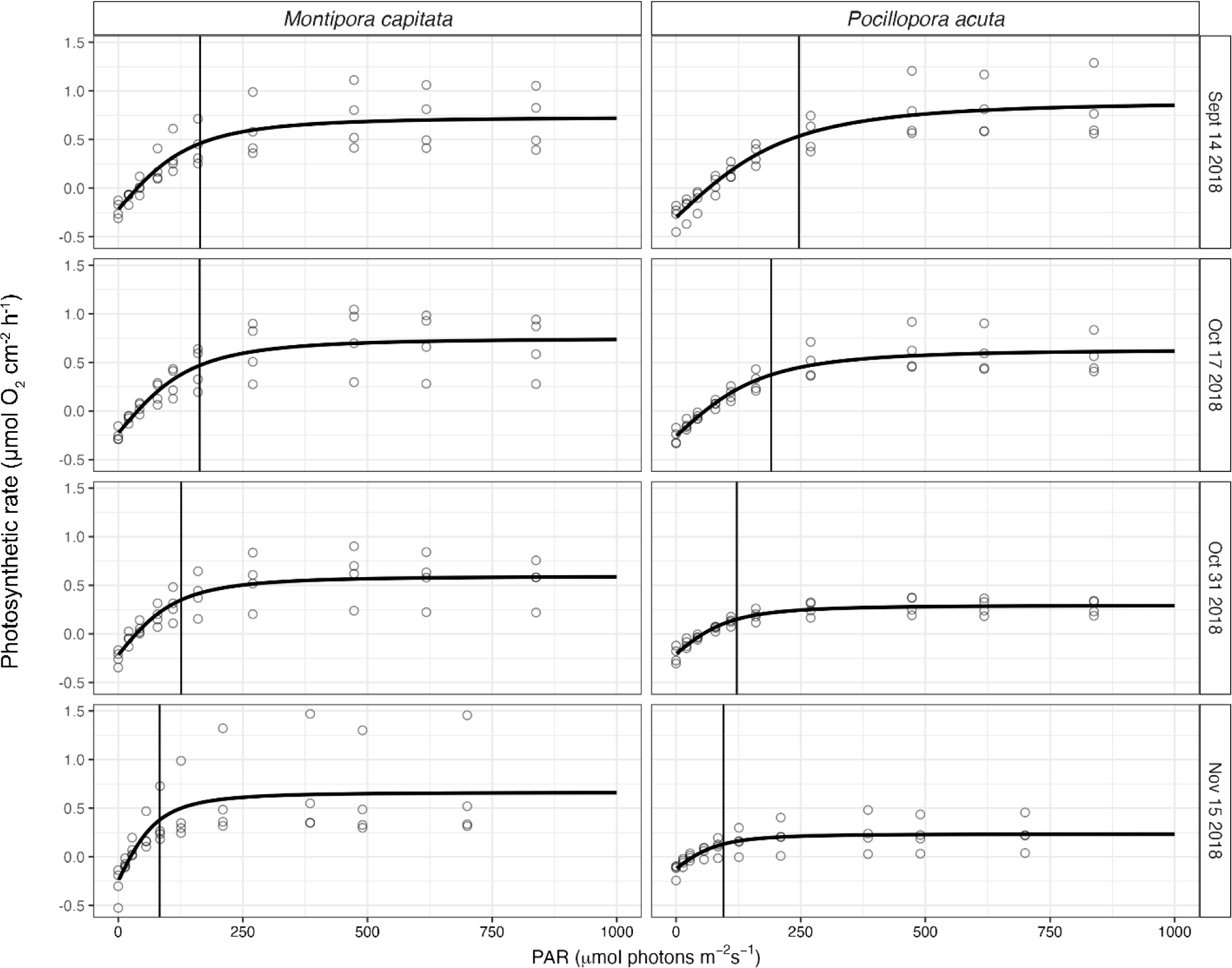
Photosynthetic Irradiance (PI) Curves for four time points during the exposure period for both species. Photosynthetic rates (μmol O_2_ cm^-2^ h^-1^) measured at a range of irradiance (μmol photons m^-2^ s^-1^) levels plotted for four time points: September 14 2018, October 17 2018, October 31 2018, and November 15 2018. PI curves were conducted to identify saturating irradiance (I_k_; solid vertical line in each plot) and ensure coral fragments were not reaching photoinhibition during respiration and photosynthesis measurements.

**Fig. S5.**
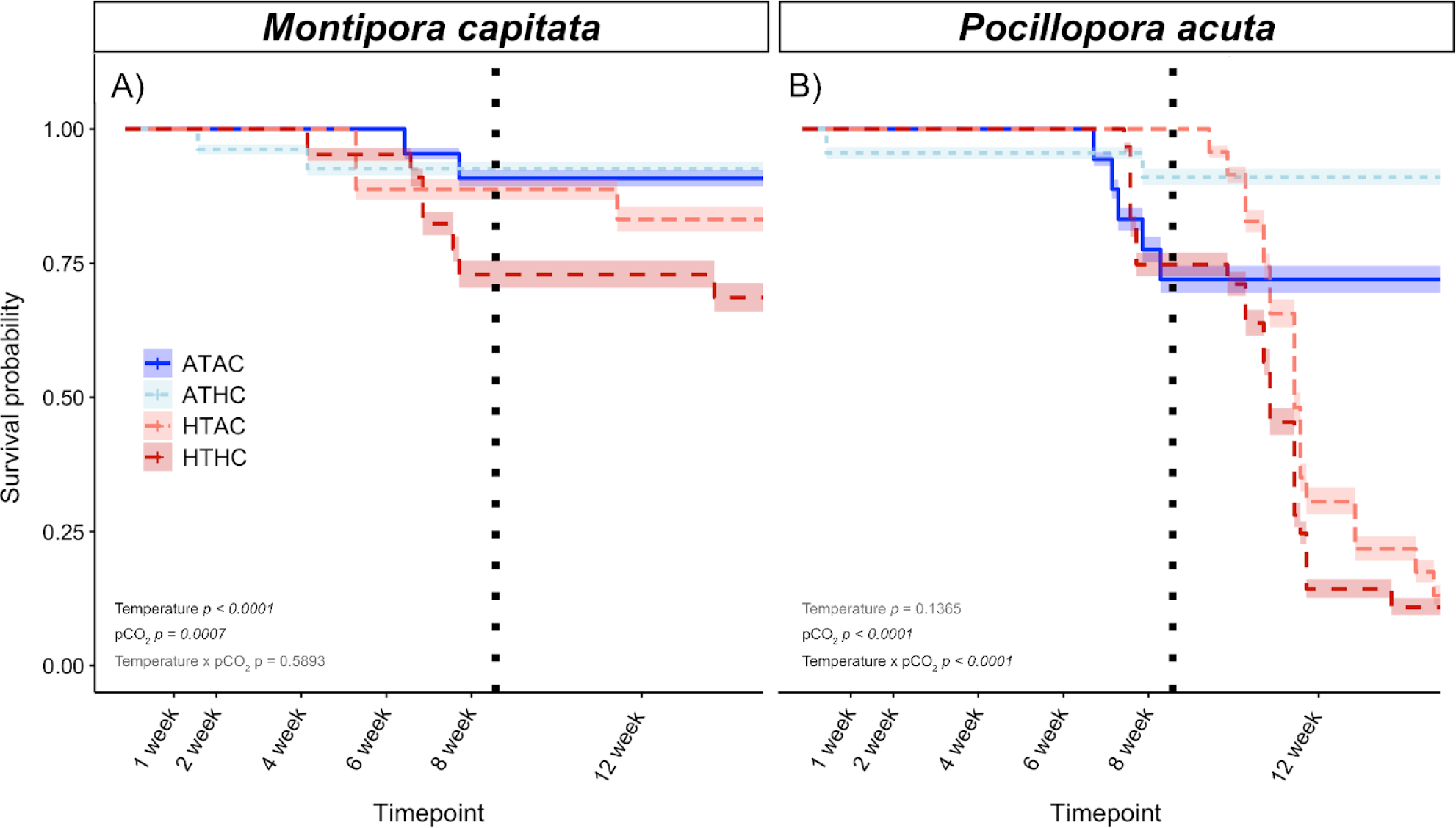
Survival probability calculated from a cox-mixed effects model for each species. Treatment is indicated by color, significant effects are in black text, and non-significant effects are in gray text. The dotted line is the point when all treatment returns to ambient level conditions. Survival probability is on a scale of 0-1.00 for both (A) *Montipora capitata* fragments. (B) *Pocillopora acuta* fragments across both exposure (day 1 - week 8) and recovery (week 8-16) periods.

**Fig. S6.**
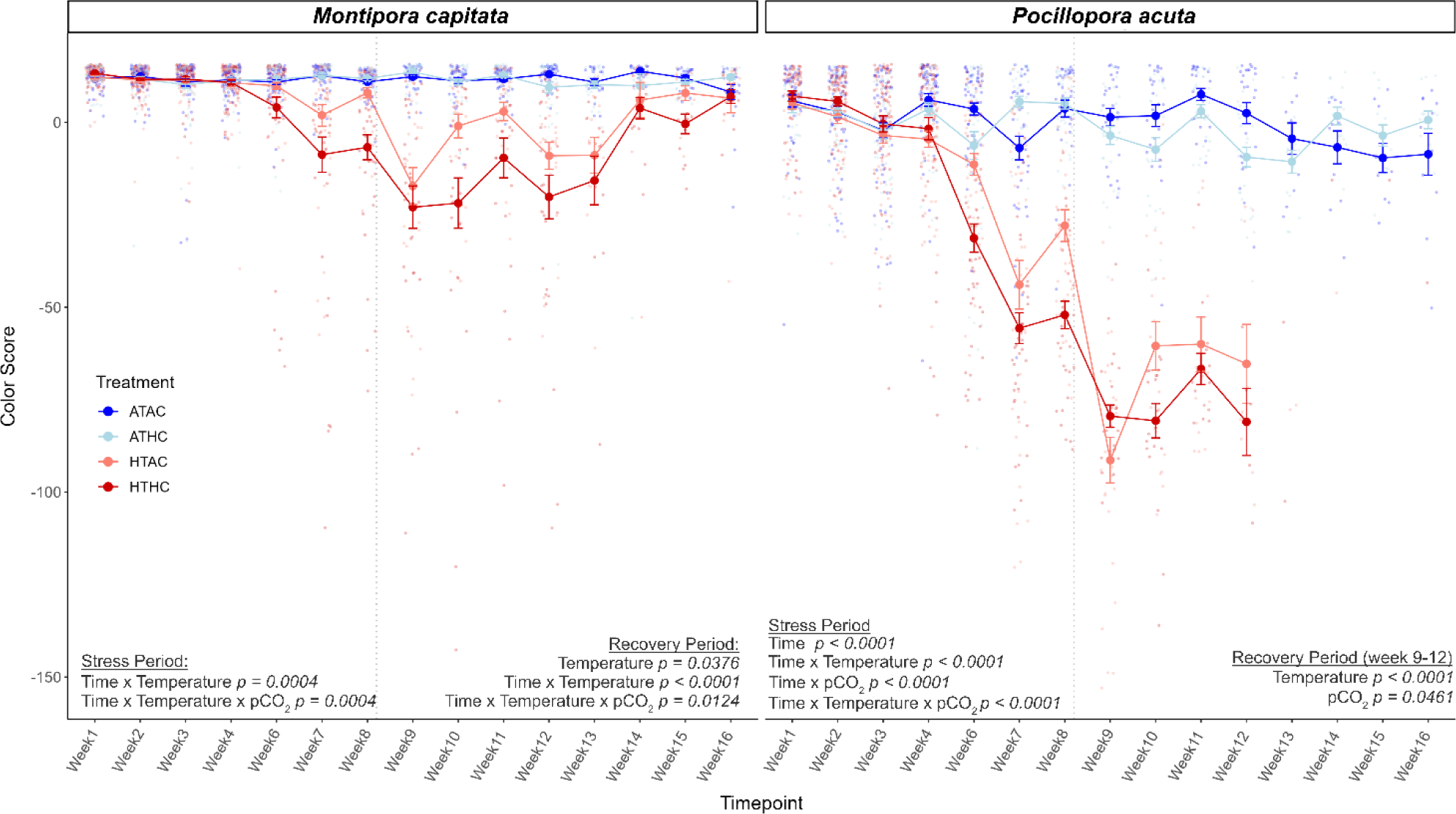
Mean ± s.e.m. weekly color score measurements through both exposure (week 1-8) and recovery (week 9-16) periods for each species. As coral bleaches, color score decreases into negative values. (A) *M. capitata* fragments. (B) *P. acuta* fragments. Significant effects from the Type III ANOVA of linear mixed model outputs are listed on each panel. Treatments are indicated by colored mean points for each time point. The dotted line is when treatment conditions return to fully ambient values (week 9).

**Fig. S7.**
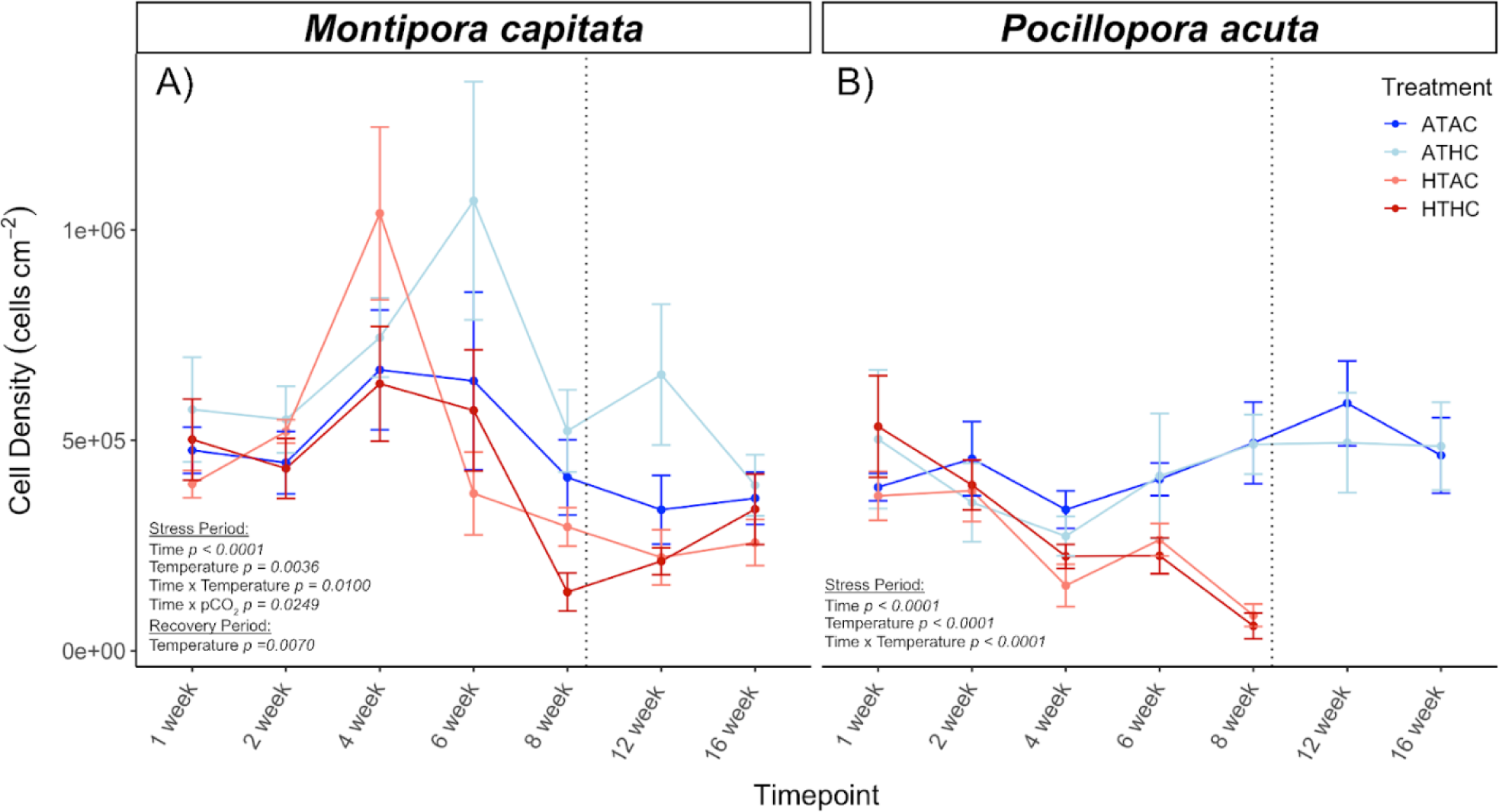
Mean endosymbiont cell density levels (cells cm^-2^) through both exposure and recovery periods for (A) *M. capitata* and (B) *P. acuta*. Treatments are indicated by color and significant effects for each species are labeled on the corresponding panel. The dotted line marks when all treatments returned to ambient level conditions at week 9. In *P. acuta*, high temperature treatments did not yield enough sample sizes to complete physiological measurements for weeks 12 and 16.

**Fig. S8.**
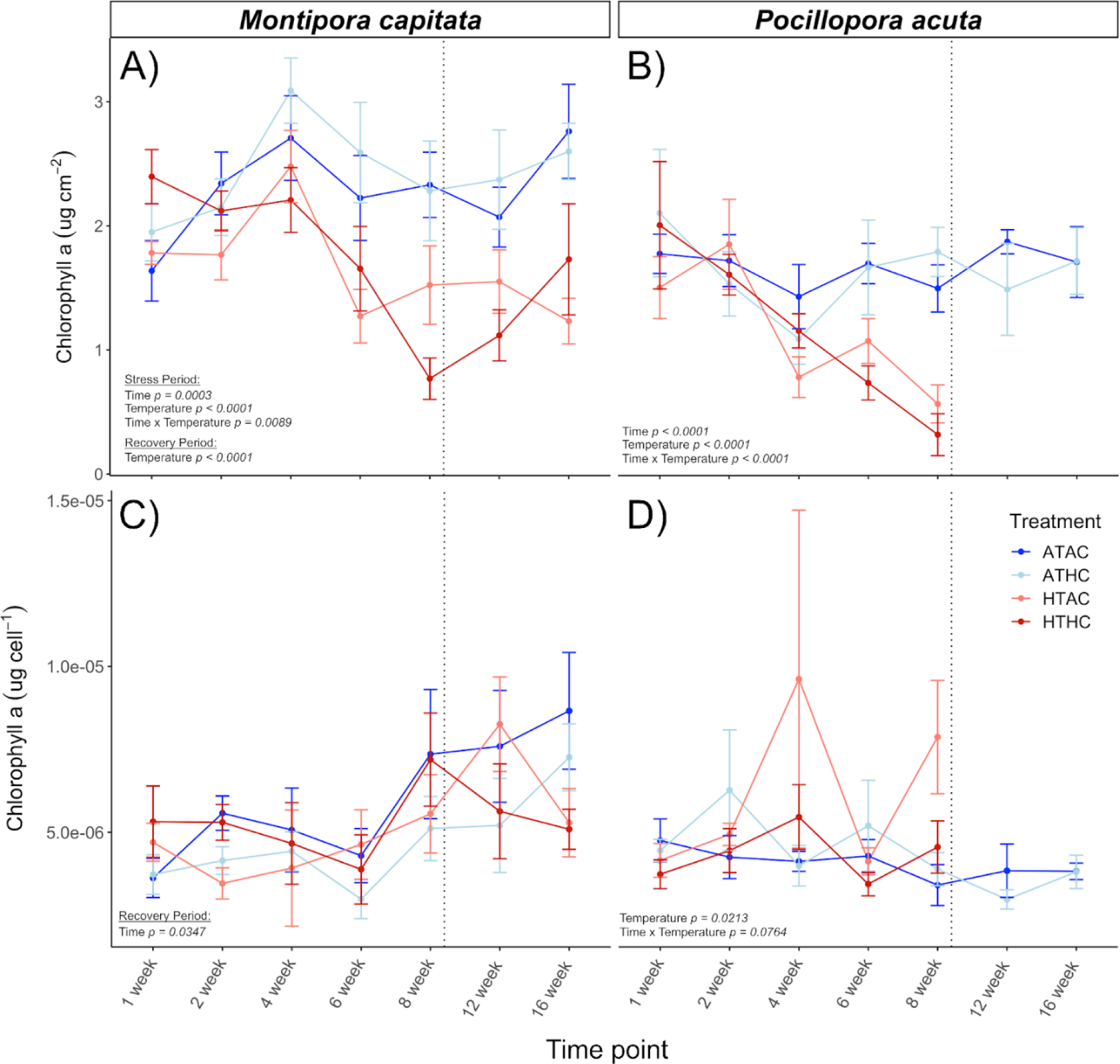
Mean chlorophyll-a-a content levels normalized to both surface area (μg cm^-2^) and endosymbiont cell density (μg cell^-1^) for (A, C) *M. capitata* and (B, D) *P. acuta*. Treatments are indicated by color and significant effects for each species are labeled on the corresponding panel. The dotted line marks when all treatments returned to ambient level conditions at week 9. In all *P. actua* panels, high temperature treatments did not yield enough sample sizes to complete physiological measurements for weeks 12 and 16.

**Fig. S9.**
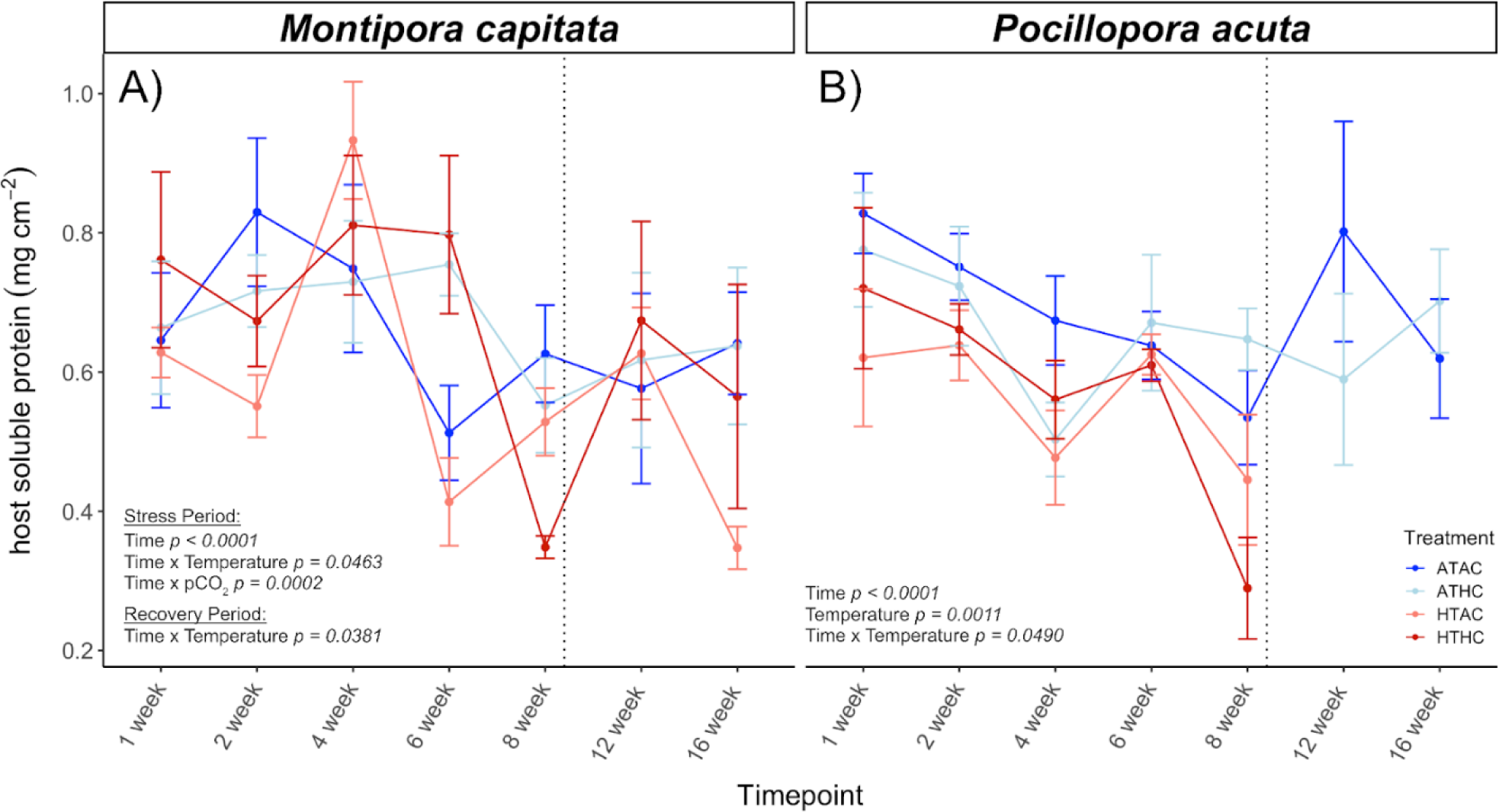
Mean host soluble protein levels normalized to surface area (mg cm^-2^) for both (A) *M. capitata* and (B) *P. acuta* in exposure and recovery periods. Treatments are indicated by color and significant effects for each species are labeled on the corresponding panel. The dotted line marks when all treatments returned to ambient level conditions at week 9. In *P. acuta*, high temperature treatments did not yield enough sample sizes to complete physiological measurements for weeks 12 and 16.

**Fig. S10.**
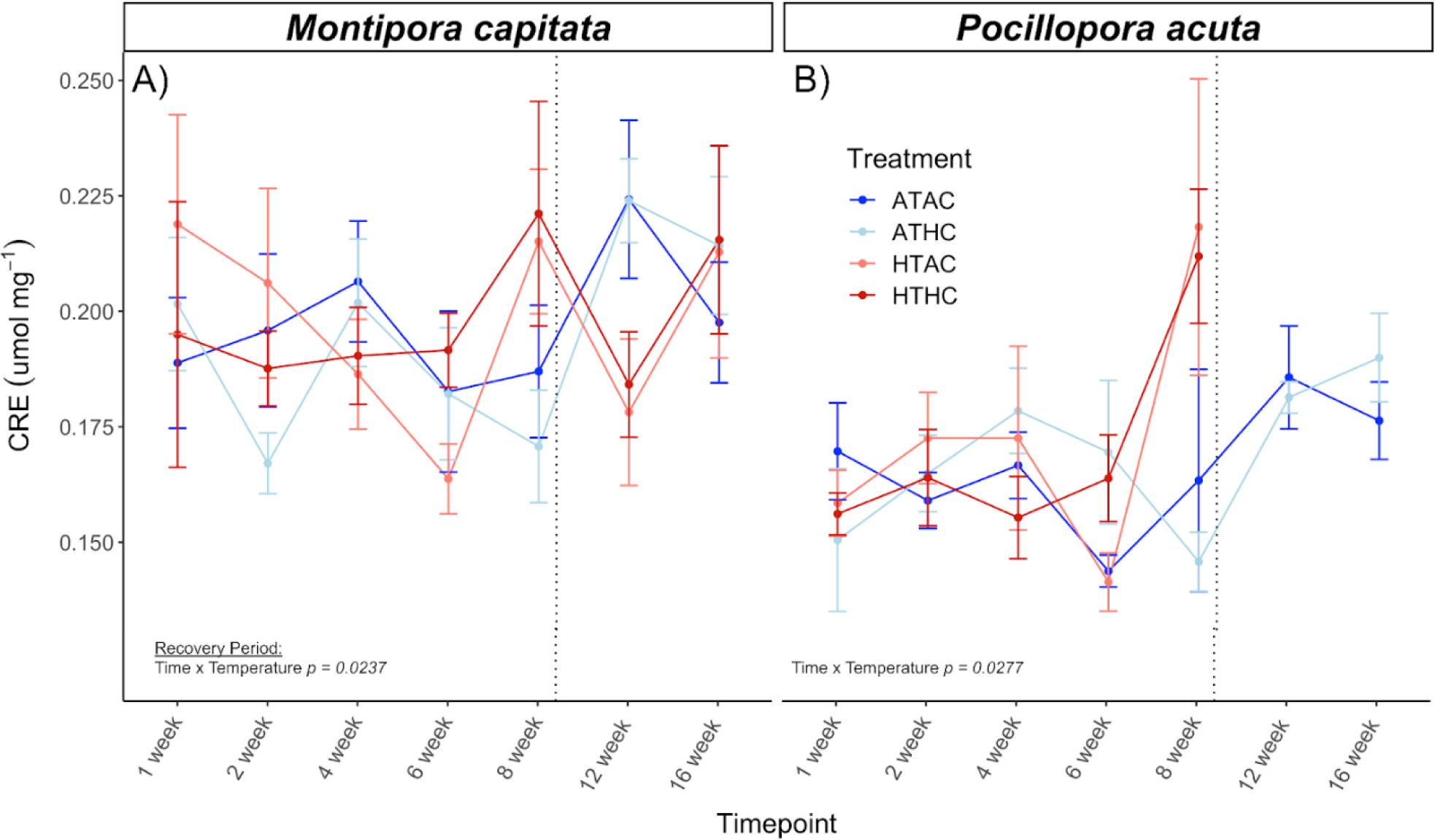
Mean host total antioxidant capacity (TAC) levels normalized to host soluble protein (CRE μmol mg^-1^) for both (A) *M. capitata* and (B) *P. acuta* through exposure and recovery periods. Treatments are indicated by color and significant effects for each species are labeled on the corresponding panel. The dotted line marks when all treatments returned to ambient level conditions at week 9. In *P. acuta*, high temperature treatments did not yield enough sample sizes to complete physiological measurements for weeks 12 and 16.

**Fig. S11.**
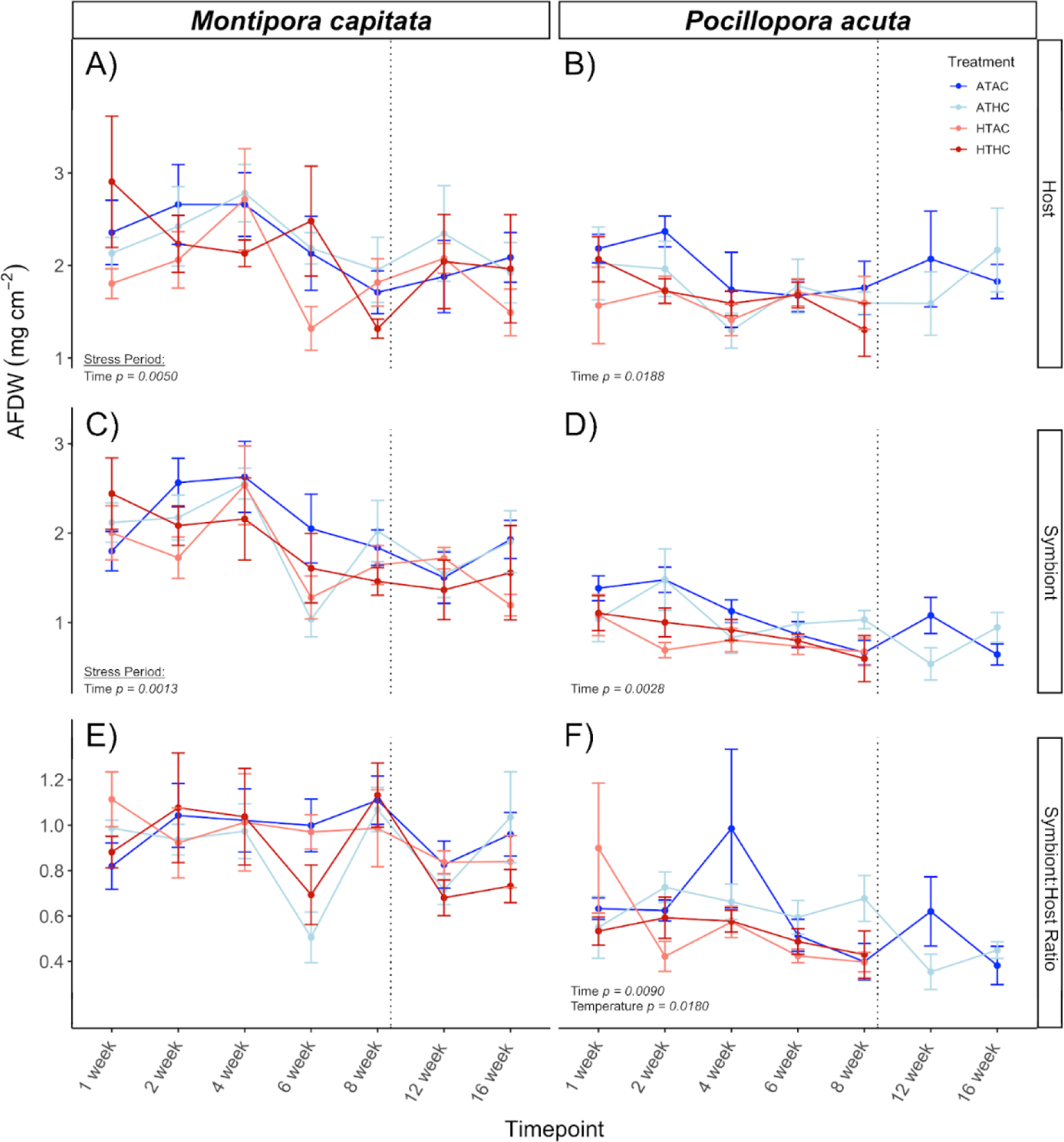
Mean tissue biomass levels measured in Ash-Free Dry Weight normalized to surface area (AFDW mg cm^-2^) for host, symbiont, and the ratio of symbiont:host for both (A, C, E) *M. capitata* and (B, D, F) *P. acuta*. Treatments are indicated by color and significant effects are labeled on the corresponding panel. The dotted line represents when treatments returned to ambient level conditions. (A,B) Host AFDW (mg cm^-2^). (C,D) Symbiont AFDW (mg cm^-2^). (E,F) Symbiont:Host AFDW Ratio. For all *P. acuta* panels, high temperature treatments did not yield enough sample sizes to complete physiological measurements for weeks 12 and 16.

**Fig. S12.**
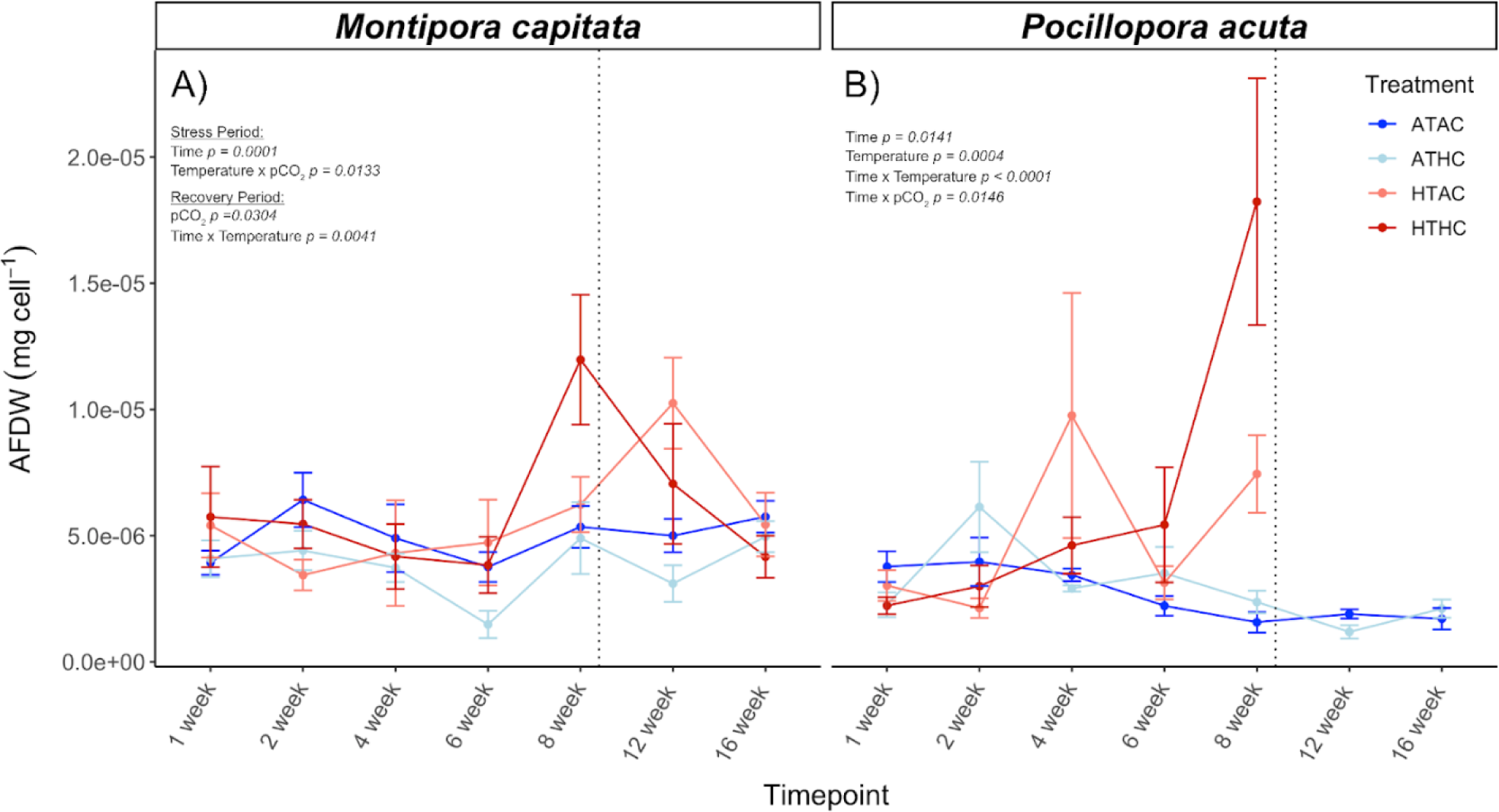
Mean symbiont cellular Ash-Free Dry Weight levels normalized to cell densities (AFDW; mg cell^-1^) for (A) *M. capitata* and (B) *P. acuta*. Treatments are indicated by color and significant effects are labeled on the corresponding panel. The dotted line represents when treatments returned to ambient level conditions. In *P. acuta*, high temperature treatments did not yield enough sample sizes to complete physiological measurements for weeks 12 and 16.

**Fig. S13.**
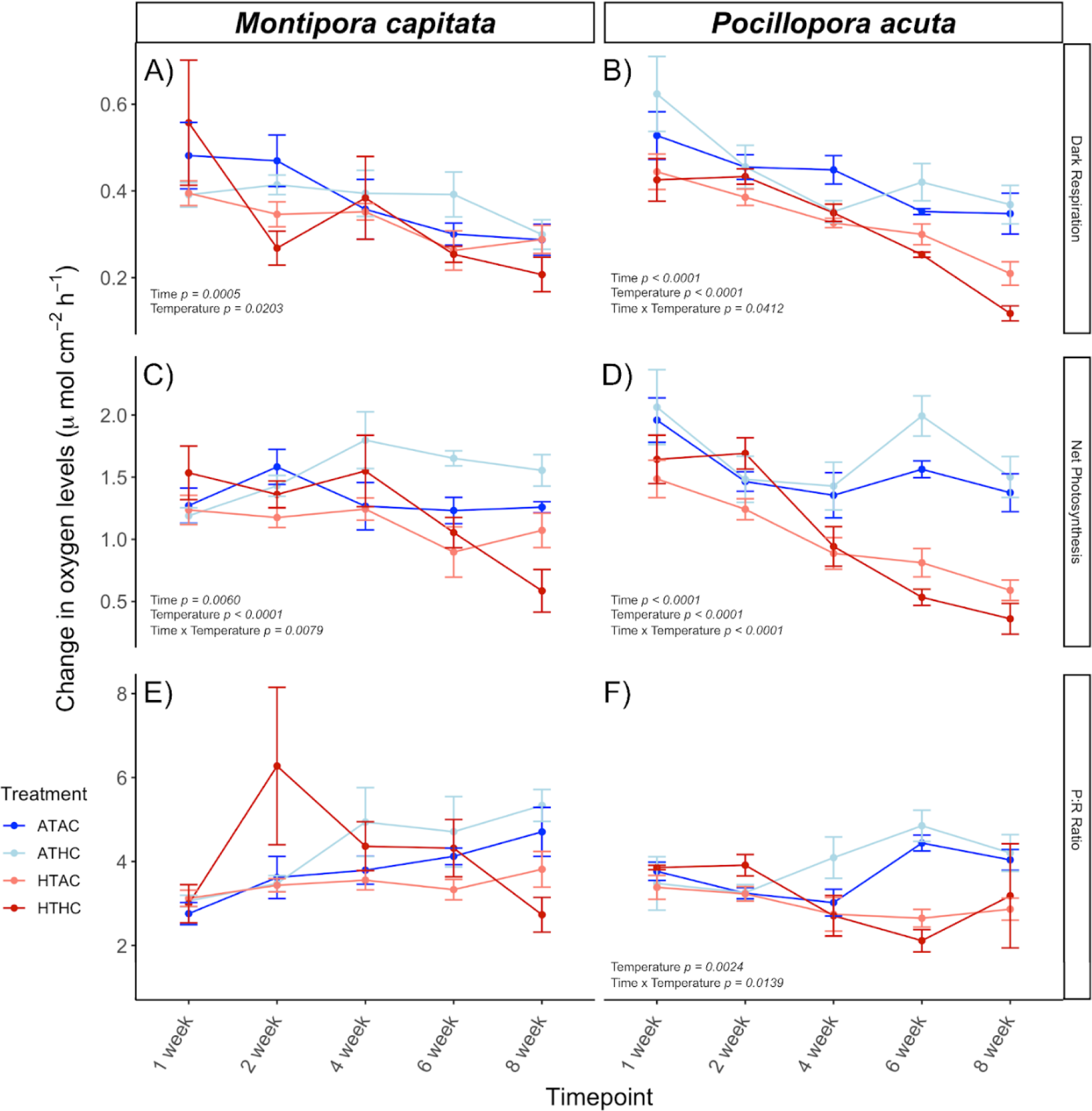
Mean dark respiration and net photosynthetic rates during the exposure period for both (A, C, E) *M. capitata* and (B, D, F) *P. acuta*. Treatments are indicated by color and significant effects are labeled on the corresponding panel. Both respiration (A,B) and photosynthetic rates (C,D) are reported in units of change in oxygen levels (μmol cm^-2^ hr^-1^). Respiration rates are the absolute value of the rate of change so that a decrease in rates is equal to lower respiration happening within the coral holobiont. Both respiration and photosynthetic rates were not measured for the recovery period. (E,F) Net photosynthetic rates (μmol cm^-2^ hr^-1^) to dark respiration rates (μmol cm^-2^ hr^-1^) ratio reported in unitless values.

**Fig. S14.**
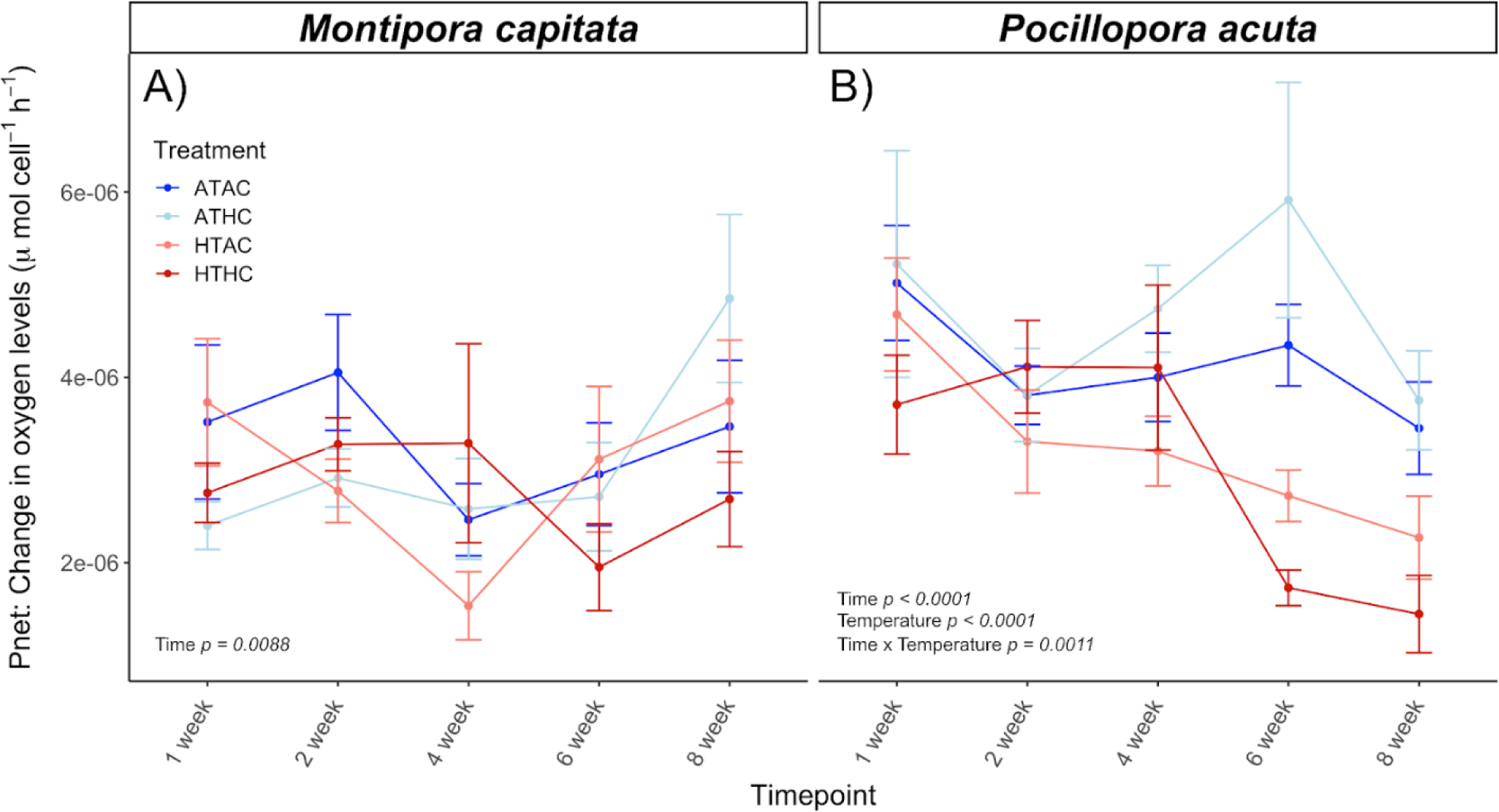
Mean cellular net photosynthetic rates measured via change in oxygen levels μmol cell^-1^ hr^-1^) and normalized to cell densities for each species (A) *M. capitata*. (B) *P. acuta*. Treatments are indicated by color and significant effects are labeled on the corresponding panel. Both respiration and photosynthetic rates were not measured for the recovery period.

**Fig. S15.**
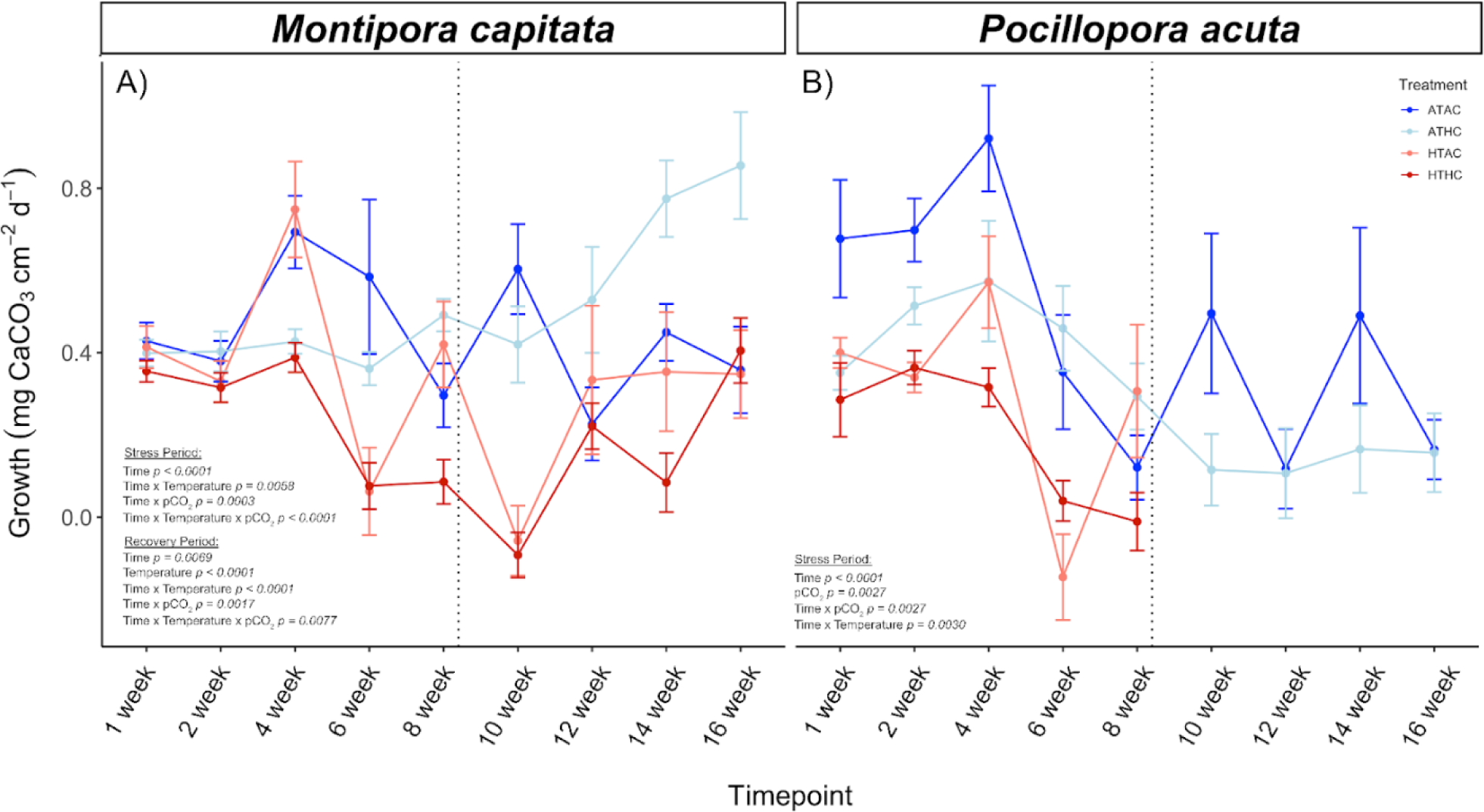
Weekly mean holobiont growth rates measured in mg CaCO_3_ cm^-2^ d^-1^ for both (A) *M. capitata* and (B) *Pocilopora acuta*. Treatments are indicated by color and significant effects are labeled on the corresponding panel. The dotted line indicates when treatment returned to ambient levels. *P. acuta* fragments did not survive in enough quantities for sufficient sample sizes during the recovery period.

**Fig. S16.**
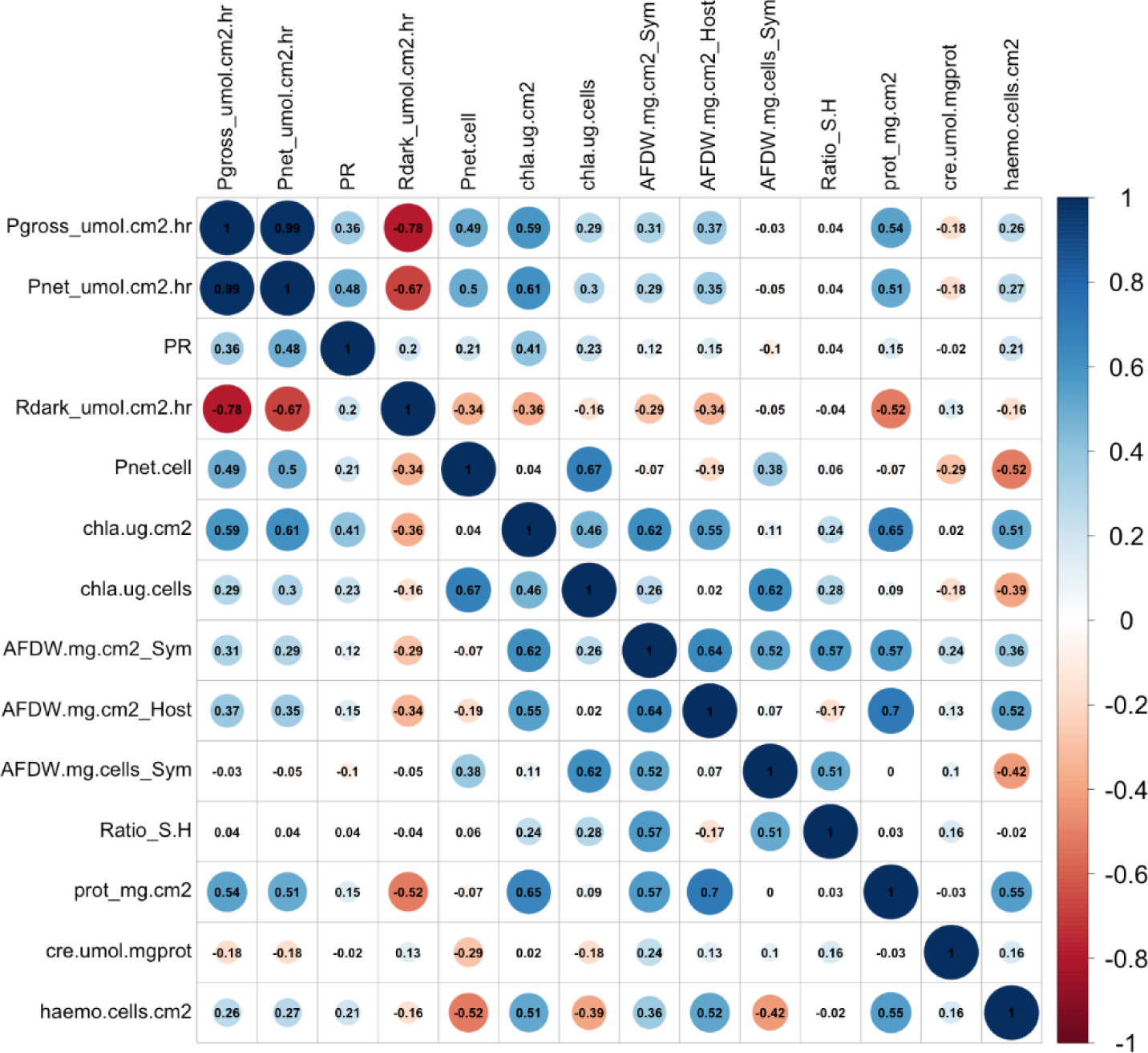
Correlation plot between all measured physiological variables (14 total). All treatments, species, and timepoints are collapsed into one figure to assess which variables are highly correlated and redundant to include in multivariate analyses. Dark blue indicates highly positively correlated (value of 1) and dark red indicates highly negatively correlated (value of -1). The size of the dot in each cell indicates the degree of correlation so that a larger dot is a higher correlation. Gross photosynthetic rates and net photosynthetic rates were highly positively correlated and therefore only net photosynthetic rates were included in multivariate analyses.

**Fig. S17.**
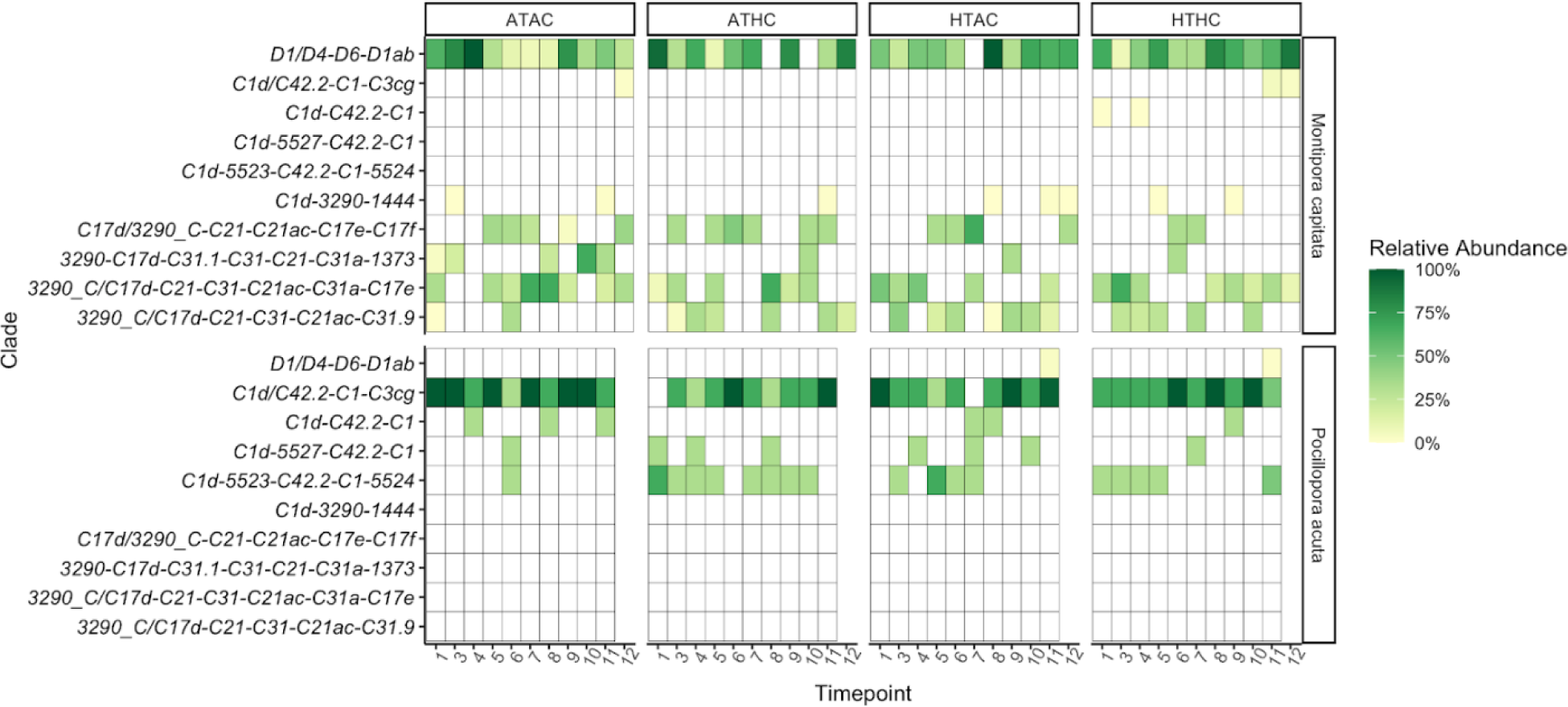
Relative abundance (%) of ITS2 Symbiodiniaceae Type Profiles. Each individual square is the average percent relative abundance per treatment per time point per species (n=3). The four column sections indicate treatments (ATAC, ATHC, HTAC, and HTHC) and the two row sections indicate host species (*Montipora capitata* and *Pocillopora acuta*). The following time points are included in the exposure period: 0 hour, 6 hour, 12 hour, 30 hour, 1, 2, 4, 6, and 8 weeks. The vertical dotted line indicates the start of the recovery period, which includes 12 and 16 weeks. *P. acuta* samples were only collected through week 12. Each square is colored by percentage relative abundance, with 100% as the darkest green and 0% as white. In the ITS2 Type Profile name, the major sequence is bolded, followed by the rest of the full type profile name.

**Fig S18.**
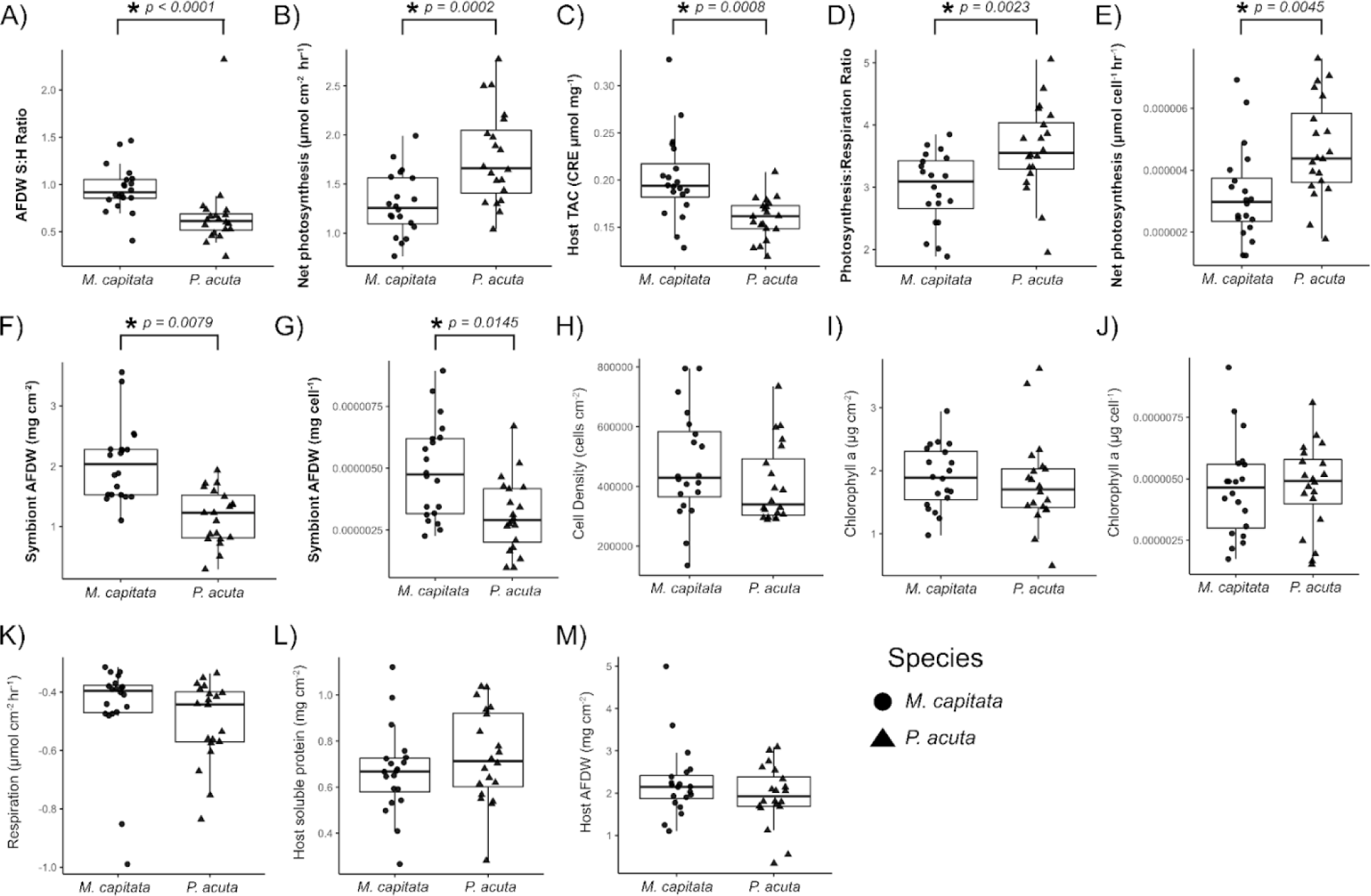
Baseline physiological measurements from week 1 ordered by most significant differences to least. (A) Ash-free dry weight symbiont:host ratio, (B) Holobiont net photosynthetic rates (μmol cm^-2^ hr^-1^), (C) Host total antioxidant capacity (CRE μmol mg^-1^), (D) Photosynthesis:Respiration ratio, (E) Cellular net photosynthetic rates (μmol cell^-1^ hr^-1^), (F) Symbiont AFDW (mg cm^-2^), (G) Symbiont cellular AFDW (mg cell^-1^), (H) Endosymbiont cell density (cells cm^-2^), (I) Holobiont chlorophyll-a-a content (μg cm^-2^), (J) Symbiont cellular chlorophyll-a-a content (μg cell^-1^), (K) Host respiration rates (μmol cm^-2^ hr^-1^), (L) Host soluble protein (mg cm^-2^), (M) Host AFDW (mg cm^-2^). *M. capitata* indicated by circles and *P. acuta* samples indicated by triangles.

**Table S1.** Total Alkalinity sample size (N), mean, and standard error (St. Error) output from the exposure period for each treatment. A linear mixed model was run to determine the effects of treatment and time point on tank conditions.

| Treatment | *N* | *Mean* | *St. Error* |
| --- | --- | --- | --- |
| ATAC | 48 | 2159.06 | 5.89 |
| ATHC | 48 | 2162.74 | 5.49 |
| HTAC | 48 | 2155.58 | 5.25 |
| HTHC | 48 | 2156.79 | 5.31 |

| Effect | *X*<sup>2</sup> | *df* | *P* |
| --- | --- | --- | --- |
| **Timepoint** | 170.0467 | 15 | **< 0.0001** |
| Treatment | 1.1034 | 3 | 0.7762 |
| Timepoint:Treatment | 43.8277 | 45 | 0.5126 |

**Table S2.** Light (PAR) output: sample size (N), mean, and standard error (St. Error) output from the exposure period for each treatment. A linear mixed model was run to determine the effects of treatment on tank conditions.

| *aov: PAR ~ Position \* Tank* |  |  |  |  |  |
| --- | --- | --- | --- | --- | --- |
| Effect | *df* | *Sum of Sq* | *Mean Sq* | *F* | *Pr(>F)* |
| Position | 5 | 23764 | 4753 | 0.155 | 0.978 |
| Tank | 11 | 498085 | 45280 | 1.475 | 0.143 |
| Position:Tank | 55 | 120265 | 2187 | 0.071 | 1.000 |

| *Light Summary Data: Stress Period* |  |  |  |
| --- | --- | --- | --- |
| Treatment | *N* | *Mean* | *St. Error* |
| ATAC | 212 | 179.70 | 9.97 |
| ATHC | 211 | 186.54 | 10.20 |
| HTAC | 210 | 176.64 | 10.84 |
| HTHC | 211 | 165.12 | 10.41 |

| *lmer: Treatment + (1 \| Date) + (1 \| Tank)* |  |  |  |
| --- | --- | --- | --- |
| Effect | *X<sup>2</sup>* | *df* | *P* |
| Treatment | 2.0687 | 3 | 0.5583 |

**Table S3.**
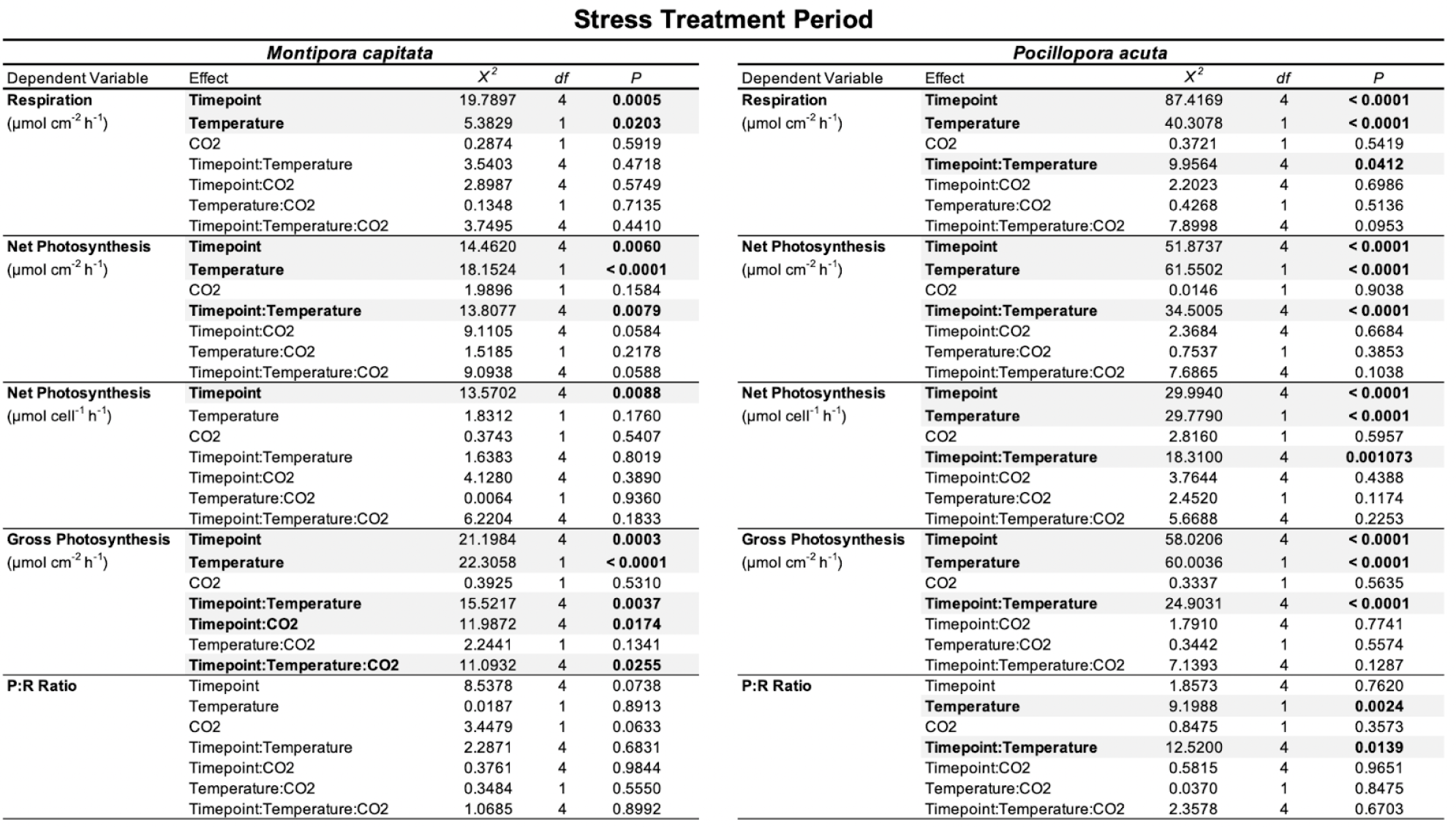
Statistical output for metabolic (photosynthetic and respiration) rates normalized to both surface area (cm^-2^) and cell density (cell^-1^). Left: *M. capitata*. Right: *P. acuta*.

**Table S4.**
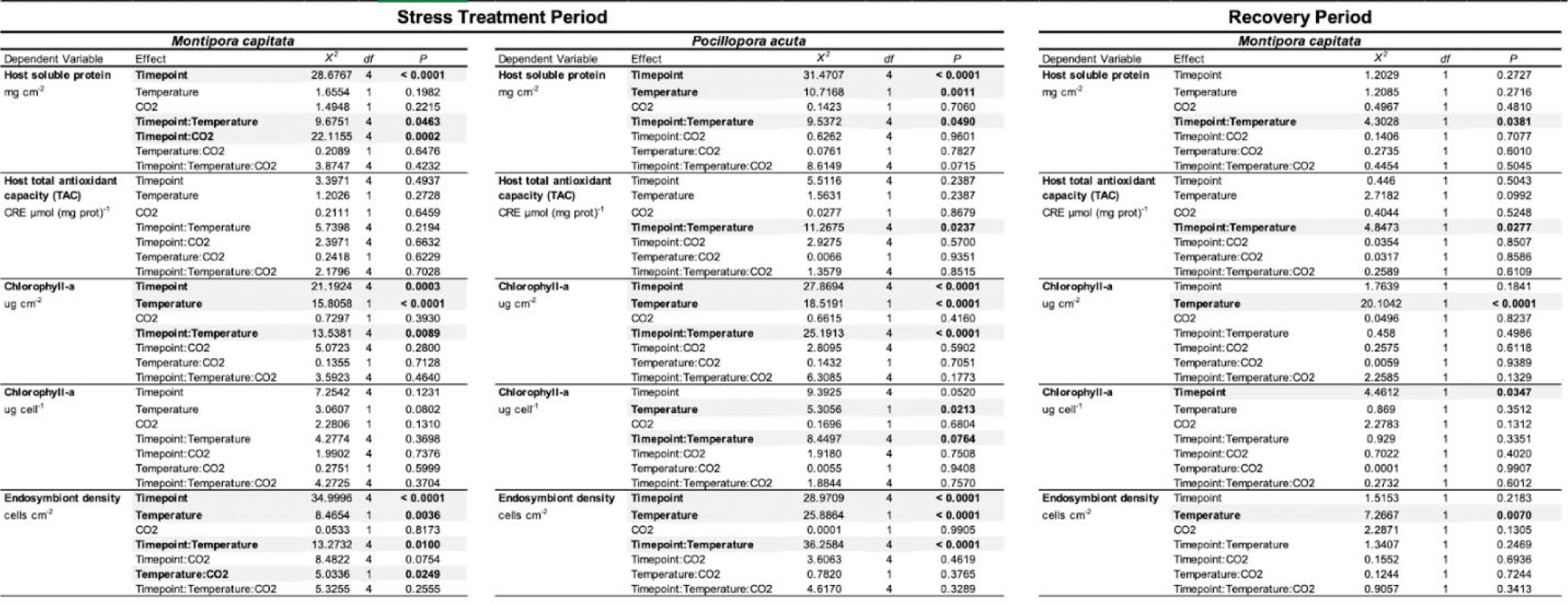
Statistical output for host soluble protein, host total antioxidant capacity, chlorophyll-a-a content, and endosymbiont densities during the exposure period for both *M. capitata* and *P. acuta* and the recovery period for *M. capitata*.

**Table S5.**
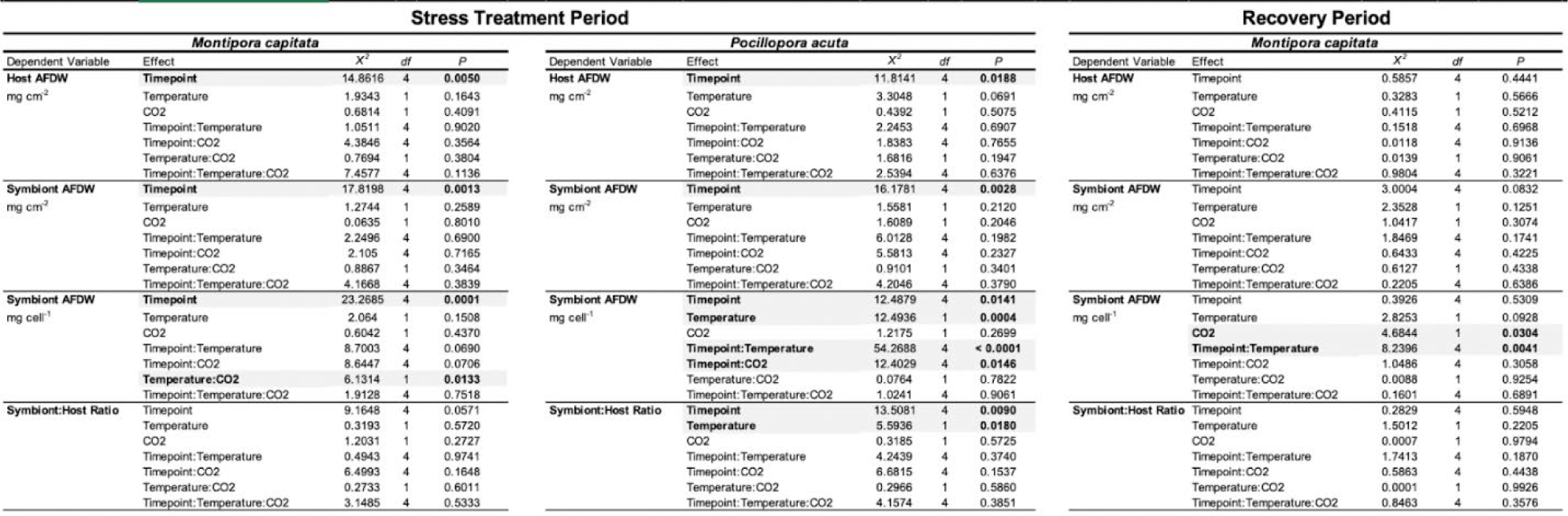
Statistical output for tissue biomass measurements: host AFDW, symbiont AFDW normalized to both surface area (cm^-2^) and cell density (cell^-1^), and symbiont:host AFDW Ratio for both species during the exposure and the recovery period for *M. capitata*.

**Table S6.**
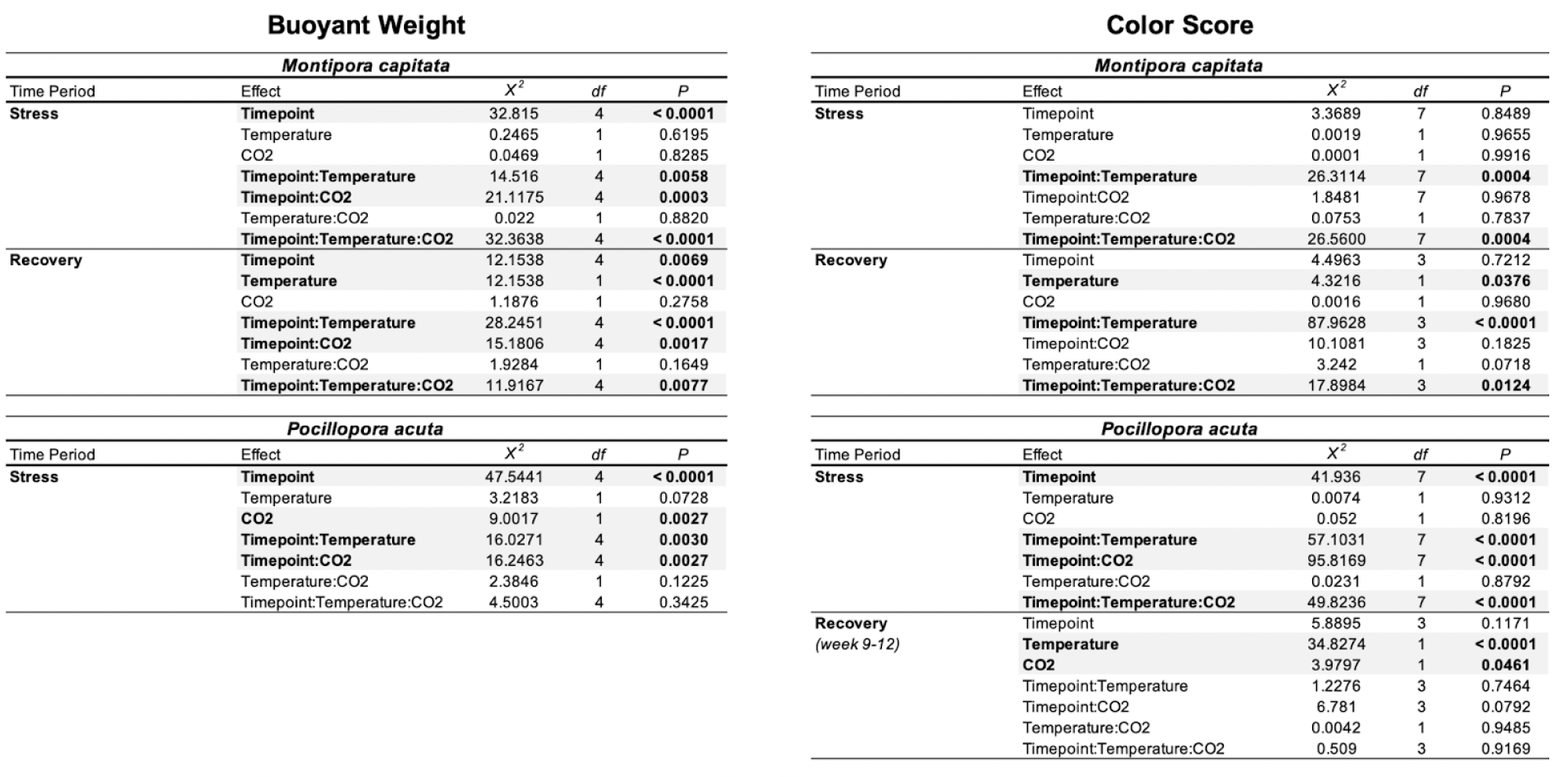
Statistical output for calcification growth rates and color score for both species during the exposure and recovery period.

**Table S7.** Statistical output from survival probability models for both *M. capitata* and *P. acuta* throughout the entire experiment.

| Type III Anova | Effect | df | $\chi^2$ | | *P* | |
| --- | --- | --- | --- | --- | --- | --- |
|  | Temperature | 1 | 21.004 |  | < 0.0001 |  |
|  | CO2 | 1 | 11.4891 |  | 0.0007 |  |
|  | Temperature:CO2 | 1 | 0.2914 |  | 0.5893 |  |
| Cox mixed-effects model fit by maximum likelihood | Effect | coef | exp(coef) | se(coef) | z | p |
| Surv(lifespan, status) ~ Temperature * CO2 +<br>(1\|PLUG.ID/Tank) | High Temperature | 1.1421189 | 3.133401 | 0.2492068 | 4.58 | < 0.0001 |
|  | High CO2 | 0.7981529 | 2.221434 | 0.2354737 | 3.39 | 0.0007 |
|  | High Temp: High CO2 | 0.1732892 | 1.18921 | 0.3210069 | 0.54 | 0.5900 |
|  |  | NULL |  | Integrated |  | Fitted |
|  | Log-likelihood | -7359.299 |  | -5183.304 |  | -4472.52 |

| Type III Anova | Effect | df | $\chi^2$ | | *P* | |
| --- | --- | --- | --- | --- | --- | --- |
|  | Temperature | 1 | 2.2173 |  | 0.1365 |  |
|  | CO2 | 1 | 40.5412 |  | < 0.0001 |  |
|  | Temperature:CO2 | 1 | 79.9506 |  | < 0.0001 |  |
| Cox mixed-effects model fit by maximum likelihood | Effect | coef | exp(coef) | se(coef) | z | p |
| Surv(lifespan, status) ~ Temperature * CO2 +<br>(1\|PLUG.ID/Tank) | High Temperature | -0.9585059 | 0.383 | 0.6436949 | -1.49 | 0.1400 |
|  | High CO2 | -4.4442835 | 0.012 | 0.6979973 | -6.37 | < 0.0001 |
|  | High Temp: High CO2 | 7.8613358 | 2594.984 | 0.8791956 | 8.94 | < 0.0001 |
|  |  | NULL |  | Integrated |  | Fitted |
|  | Log-likelihood | -24130.77 |  | NaN |  | -18100.5 |

**Table S8.**
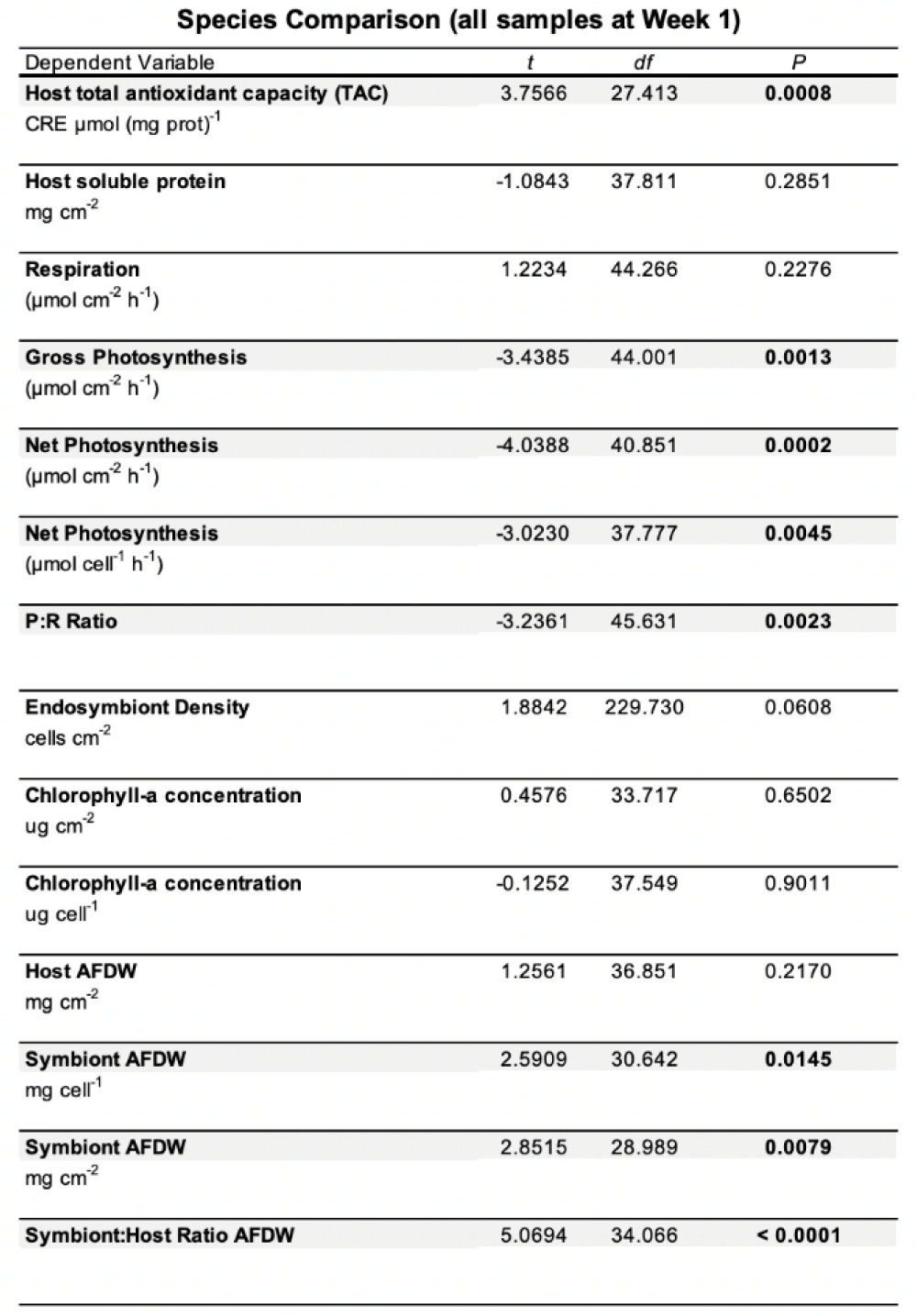
Statistical output for baseline physiological measurement comparison between *M. capitata* and *P. acuta* fragments at the initial week 1 measurement.

| Species Comparison (all samples at Week 1) |  |  |  |
| --- | --- | --- | --- |
| Dependent Variable | *t* | *df* | *P* |
| **Host total antioxidant capacity (TAC)**<br>CRE $\mu\text{mol (mg prot)}^{-1}$ | 3.7566 | 27.413 | **0.0008** |
| **Host soluble protein**<br>mg $\text{cm}^{-2}$ | -1.0843 | 37.811 | 0.2851 |
| **Respiration**<br>($\mu\text{mol cm}^{-2} \text{ h}^{-1}$) | 1.2234 | 44.266 | 0.2276 |
| **Gross Photosynthesis**<br>($\mu\text{mol cm}^{-2} \text{ h}^{-1}$) | -3.4385 | 44.001 | **0.0013** |
| **Net Photosynthesis**<br>($\mu\text{mol cm}^{-2} \text{ h}^{-1}$) | -4.0388 | 40.851 | **0.0002** |
| **Net Photosynthesis**<br>($\mu\text{mol cell}^{-1} \text{ h}^{-1}$) | -3.0230 | 37.777 | **0.0045** |
| **P:R Ratio** | -3.2361 | 45.631 | **0.0023** |
| **Endosymbiont Density**<br>cells $\text{cm}^{-2}$ | 1.8842 | 229.730 | 0.0608 |
| **Chlorophyll-a concentration**<br>ug $\text{cm}^{-2}$ | 0.4576 | 33.717 | 0.6502 |
| **Chlorophyll-a concentration**<br>ug $\text{cell}^{-1}$ | -0.1252 | 37.549 | 0.9011 |
| **Host AFDW**<br>mg $\text{cm}^{-2}$ | 1.2561 | 36.851 | 0.2170 |
| **Symbiont AFDW**<br>mg $\text{cell}^{-1}$ | 2.5909 | 30.642 | **0.0145** |
| **Symbiont AFDW**<br>mg $\text{cm}^{-2}$ | 2.8515 | 28.989 | **0.0079** |
| **Symbiont:Host Ratio AFDW** | 5.0694 | 34.066 | **< 0.0001** |

**Table S9.** Statistical output from a permutational multivariate ANOVA (PERMANOVA) on the multivariate physiology states for each species during the exposure and the recovery period for *M. capitata*.

STRESS PERIOD
| **PERMANOVA: Timepoint \* Temperature \* CO2 \* Species** |  |  |  |  |  |
| --- | --- | --- | --- | --- | --- |
| Effect | df | Sum of Squares | R2 | F | Pr(>F) |
| Timepoint | 4 | 203.56 | 0.07022 | 5.4788 | **0.001** |
| Temperature | 1 | 130.16 | 0.04490 | 14.0127 | **0.001** |
| CO2 | 1 | 13.74 | 0.00474 | 1.4789 | 0.166 |
| Species | 1 | 298.93 | 0.10311 | 32.1831 | **0.001** |
| Timepoint:Temperature | 4 | 115.98 | 0.04001 | 3.1218 | **0.001** |
| Timepoint:CO2 | 4 | 69.68 | 0.02404 | 1.8755 | **0.008** |
| Temperature:CO2 | 1 | 9.13 | 0.00315 | 0.9830 | 0.403 |
| Timepoint:Species | 4 | 145.09 | 0.05005 | 3.9051 | **0.001** |
| Temperature:Species | 1 | 21.56 | 0.00744 | 2.3213 | **0.029** |
| CO2:Species | 1 | 13.25 | 0.00457 | 1.4270 | 0.209 |
| Timepoint:Temperature:CO2 | 4 | 62.88 | 0.02169 | 1.6926 | **0.033** |
| Timepoint:Temperature:Species | 4 | 32.22 | 0.01111 | 0.8672 | 0.652 |
| Timepoint:CO2:Species | 4 | 36.75 | 0.01268 | 0.9892 | 0.456 |
| Temperature:CO2:Species | 1 | 3.11 | 0.00107 | 0.3350 | 0.936 |
| Timepoint:Temperature:CO2:Species | 4 | 33.88 | 0.01169 | 0.9119 | 0.545 |

| **Montipora capitata PERMANOVA: Timepoint \* Temperature \* CO2** |  |  |  |  |  |
| --- | --- | --- | --- | --- | --- |
| Effect | df | Sum of Squares | R2 | F | Pr(>F) |
| Timepoint | 4 | 184.65 | 0.12682 | 4.2291 | **0.001** |
| Temperature | 1 | 39.29 | 0.02699 | 3.5995 | **0.005** |
| CO2 | 1 | 20.27 | 0.01393 | 1.8574 | 0.091 |
| Timepoint:Temperature | 4 | 60.73 | 0.04171 | 1.3909 | 0.099 |
| Timepoint:CO2 | 4 | 71.81 | 0.04932 | 1.6447 | **0.036** |
| Temperature:CO2 | 1 | 5.78 | 0.00397 | 0.5298 | 0.792 |
| Timepoint:Temperature:CO2 | 4 | 58.28 | 0.04003 | 1.3349 | 0.135 |

| **Pocillopora acuta PERMANOVA: Timepoint \* Temperature \* CO2** |  |  |  |  |  |
| --- | --- | --- | --- | --- | --- |
| Effect | df | Sum of Squares | R2 | F | Pr(>F) |
| Timepoint | 4 | 171.63 | 0.12002 | 4.3344 | **0.001** |
| Temperature | 1 | 139.81 | 0.09777 | 14.1231 | **0.001** |
| CO2 | 1 | 4.67 | 0.00326 | 0.4716 | 0.824 |
| Timepoint:Temperature | 4 | 117.25 | 0.082 | 2.9611 | **0.001** |
| Timepoint:CO2 | 4 | 42.69 | 0.02985 | 1.0781 | 0.377 |
| Temperature:CO2 | 1 | 8.25 | 0.00577 | 0.8334 | 0.487 |
| Timepoint:Temperature:CO2 | 4 | 44.86 | 0.03137 | 1.133 | 0.314 |

| **Montipora capitata PERMANOVA: Timepoint \* Temperature \* CO2** |  |  |  |  |  |
| --- | --- | --- | --- | --- | --- |
| Effect | df | Sum of Squares | R2 | F | Pr(>F) |
| Timepoint | 4 | 40.95 | 0.0711 | 2.5587 | **0.013** |
| Temperature | 1 | 46.70 | 0.0811 | 5.8358 | **0.002** |
| CO2 | 1 | 5.56 | 0.0097 | 0.6945 | 0.621 |
| Timepoint:Temperature | 4 | 29.98 | 0.0521 | 1.8732 | 0.060 |
| Timepoint:CO2 | 4 | 14.18 | 0.0246 | 0.8861 | 0.507 |
| Temperature:CO2 | 1 | 2.39 | 0.0042 | 0.2987 | 0.921 |
| Timepoint:Temperature:CO2 | 4 | 12.08 | 0.0210 | 0.7550 | 0.661 |

**Table S10.** Statistical output from a permutational multivariate ANOVA (PERMANOVA) on the ITS2 full type profiles for each species for the entire time series.

ITS2
| **PERMANOVA: Timepoint \* Temperature \* CO2 \* Species** |  |  |  |  |  |
| --- | --- | --- | --- | --- | --- |
| Effect | df | Sum of Squares | R2 | F | Pr(>F) |
| **Timepoint** | 10 | 119.18 | 0.04767 | 1.3438 | **0.023** |
| Temperature | 1 | 4.44 | 0.00178 | 0.5008 | 0.898 |
| CO2 | 1 | 9.38 | 0.00375 | 1.0575 | 0.394 |
| **Species** | 1 | 329.71 | 0.13188 | 37.1762 | **0.001** |
| Timepoint:Temperature | 10 | 92.17 | 0.03687 | 1.0392 | 0.384 |
| Timepoint:CO2 | 10 | 81.46 | 0.03258 | 0.9184 | 0.699 |
| Temperature:CO2 | 1 | 13.62 | 0.00545 | 1.5361 | 0.121 |
| Timepoint:Species | 10 | 95.53 | 0.03821 | 1.1968 | 0.119 |
| Temperature:Species | 1 | 4.52 | 0.00181 | 0.5100 | 0.859 |
| CO2:Species | 1 | 9.32 | 0.00373 | 1.0505 | 0.407 |
| Timepoint:Temperature:CO2 | 10 | 45.77 | 0.01831 | 0.5160 | 1.000 |
| Timepoint:Temperature:Species | 10 | 88.20 | 0.03528 | 1.1050 | 0.230 |
| Timepoint:CO2:Species | 10 | 69.92 | 0.02797 | 0.8760 | 0.776 |
| Temperature:CO2:Species | 1 | 13.35 | 0.00534 | 1.5057 | 0.138 |
| Timepoint:Temperature:CO2:Species | 10 | 42.32 | 0.01693 | 0.5302 | 1.000 |

| **Montipora capitata PERMANOVA: Timepoint \* Temperature \* CO2** |  |  |  |  |  |
| --- | --- | --- | --- | --- | --- |
| Effect | df | Sum of Squares | R2 | F | Pr(>F) |
| **Timepoint** | 10 | 107.80 | 0.10286 | 1.3377 | **0.030** |
| Temperature | 1 | 7.88 | 0.00752 | 0.9782 | 0.457 |
| CO2 | 1 | 6.37 | 0.00608 | 0.7903 | 0.593 |
| Timepoint:Temperature | 10 | 94.42 | 0.09010 | 1.1717 | 0.161 |
| Timepoint:CO2 | 10 | 68.68 | 0.06554 | 0.8523 | 0.810 |
| Temperature:CO2 | 1 | 10.94 | 0.01044 | 1.3573 | 0.215 |
| Timepoint:Temperature:CO2 | 10 | 42.76 | 0.04080 | 0.5306 | 1.000 |

| **Pocillopora acuta PERMANOVA: Timepoint \* Temperature \* CO2** |  |  |  |  |  |
| --- | --- | --- | --- | --- | --- |
| Effect | df | Sum of Squares | R2 | F | Pr(>F) |
| Timepoint | 10 | 44.00 | 0.07458 | 0.9719 | 0.501 |
| Temperature | 1 | 2.25 | 0.00381 | 0.4469 | 0.795 |
| CO2 | 1 | 7.63 | 0.01292 | 1.5159 | 0.191 |
| Timepoint:Temperature | 10 | 55.42 | 0.09394 | 1.2242 | 0.177 |
| Timepoint:CO2 | 10 | 40.50 | 0.06865 | 0.8946 | 0.630 |
| Temperature:CO2 | 1 | 10.59 | 0.01795 | 2.1055 | 0.068 |
| Timepoint:Temperature:CO2 | 10 | 32.21 | 0.05460 | 0.7115 | 0.916 |

**Table S11.** Photosynthetic curve output (mean ± standard error, n=4) for maximal photosynthetic rate (Am; µmol O_2_ cm^-2^ h^-1^), apparent quantum yield (AQY, ф; mol O_2_ mol photons^-1^), Compensation irradiance (IC; PAR), saturating irradiance (I_k_; PAR), and light enhanced dark respiration rate (Rd; µmol O_2_ cm^-2^ h^-1^).c

| Species | Metric | Sept 18 2018 | Oct 17 2018 | Oct 31 2018 | Nov 15 2018 |
| --- | --- | --- | --- | --- | --- |
| *Montipora capitata* | Am | 0.95 ± 0.196 | 0.98 ± 0.178 | 0.81 ± 0.150 | 0.91 ± 0.328 |
|  | AQY | 0.01 ± 0.001 | 0.01 ± 0.001 | 0.01 ± 0.001 | 0.01 ± 0.003 |
|  | Ic | 40.40 ± 5.467 | 40.96 ± 5.092 | 38.02 ± 5.459 | 25.38 ± 2.962 |
|  | Ik | 164.08 ± 14.970 | 163.35 ± 22.658 | 126.51 ± 6.308 | 83.19 ± 7.270 |
|  | Rd | 0.22 ± 0.041 | 0.23 ± 0.028 | 0.22 ± 0.034 | 0.25 ± 0.066 |
| *Pocillopora acuta* | Am | 1.19 ± 0.257 | 0.89 ± 0.144 | 0.50 ± 0.079 | 0.36 ± 0.120 |
|  | AQY | 0.005 ± 0.001 | 0.005 ± 0.001 | 0.004 ± 0.001 | 0.004 ± 0.001 |
|  | Ic | 64.40 ± 8.240 | 58.28 ± 4.172 | 54.16 ± 4.208 | 50.12 ± 20.932 |
|  | Ik | 246.29 ± 26.716 | 190.53 ± 9.757 | 121.90 ± 5.651 | 95.21 ± 7.303 |
|  | Rd | 0.30 ± 0.069 | 0.26 ± 0.038 | 0.21 ± 0.040 | 0.12 ± 0.031 |

## Notes

https://github.com/hputnam/Acclim_Dynamics

### Competing Interest Statement

The authors have declared no competing interest.

### Summary of Updates

Updated introduction, results, and discussion text, figures, and captions.

